# PARP14 inhibition restores PD-1 immune checkpoint inhibitor response following IFNγ-driven adaptive resistance

**DOI:** 10.1101/2022.11.18.517143

**Authors:** Chun Wai Wong, Christos Evangelou, Kieran N. Sefton, Rotem Leshem, Kleita Sergiou, Macarena Lucia Fernandez Carro, Erez Uzuner, Holly Mole, Brian A. Telfer, Daniel J. Wilcock, Michael P. Smith, Kaiko Kunii, Nicholas R. Perl, Paul Lorigan, Kaye J. Williams, Patricia E. Rao, Raghavendar T. Nagaraju, Mario Niepel, Adam F.L. Hurlstone

## Abstract

Adaptive resistance limits immune checkpoint blockade therapy (ICBT) response duration and magnitude. Interferon γ (IFNγ), a critical cytokine that promotes cellular immunity, also induces adaptive resistance to ICBT. Using syngeneic mouse tumour models, we confirmed that chronic IFNγ exposure confers resistance to anti-Programmed cell death protein 1 (α-PD-1) therapy. We identified consistent upregulation of poly-ADP ribosyl polymerase 14 (PARP14) in both chronic IFNγ-treated cancer cells and patient melanoma with elevated *IFNG* expression. Knockdown or pharmacological inhibition of PARP14 increased effector T cell infiltration into tumours derived from cells pre-treated with IFNγ and decreased the presence of regulatory T cells, leading to restoration of α-PD-1 sensitivity. Finally, we determined that tumours which spontaneously relapsed following α-PD-1 therapy could be re-sensitised upon receiving PARP14 inhibitor treatment, establishing PARP14 as an actionable target to reverse IFNγ-driven ICBT resistance.

## Introduction

Programmed cell death protein 1 (PD-1) is an immune checkpoint protein highly expressed on tumour-infiltrating lymphocytes. Interaction with its ligand, programmed death ligand-1 (PD-L1), on cells in the tumour microenvironment (TME) and tumour draining lymph nodes promotes tumour immune evasion—and thereby disease progression—by suppressing effector T cell proliferation, migration, cytotoxicity, and anti-tumour immune responses^1^. Therapeutics impeding immune checkpoint interactions have revolutionised treatment of melanoma and other solid cancers such as non-small cell lung cancer, bladder carcinoma, and microsatellite instability-high (MSI-H) cancers^2^. In melanoma, PD-1 blockade reverses T cell exhaustion and increases central memory CD4^+^ T cell levels, improving overall T cell response through enhanced activation and cytotoxicity^3^. However, the effectiveness of immune checkpoint blockade therapy (ICBT) is limited by multiple drug resistance mechanisms. While primary resistance is widespread, cases where tumours initially respond but subsequently relapse within months or years are also common, with many such tumours exhibiting evidence of immunoediting and increased expression of immune checkpoint molecules^4,5^. The resulting restoration of an immunosuppressive TME leads once more to T cell exhaustion, impeding tumour cell clearance^6^.

Mechanisms of ICBT resistance are incompletely understood. As a key component of the inflammatory milieu that characterises the TME, the cytokine interferon γ (IFNγ) exerts divergent effects on tumour immune responses and tumour progression, as well as response to ICBT. Its role in tumour immunosurveillance is well established^7^, and targets of IFNγ signalling are robust biomarkers of clinical response in ICBT^8^. Moreover, defects in genes implicated in the IFNγ pathway are enriched in tumours displaying ICBT resistance^9,10^. Conversely, though, elevated IFNγ at tumour sites has been implicated in immune evasion and ICBT resistance^5^. Furthermore, tumours derived from cells treated with IFNγ prior to implantation in syngeneic mice are resistant to ICBT^11^; while *in vivo* CRISPR screens revealed IFNγ signalling as a driver of ICBT resistance in multiple syngeneic mouse tumour implantation models^12^. The upregulation of MHC and antigen-processing factors by the transcription factor Signal transducer and activator of transcription 1 (STAT1) downstream of IFNγ augments tumour antigenicity and thereby increases tumour cell recognition by T effector cells albeit at the cost of inhibiting NK cell responses; in contrast, the duration and strength of anti-tumour responses are impeded by IFNγ-induced immunomodulatory molecules, including PD-L1, which confer immune homeostasis^13^. In addition, induction of IRF2, a STAT1 target gene product, in T cells alos results in interferon-mediated T-cell exhaustion in multiple tumour types^14^. The identification of actionable targets mediating IFNγ-driven adaptive resistance is urgently needed to improve the success of ICBT.

In this study, we investigated IFNγ-driven reprogramming of gene expression in tumour cells associated with adaptive resistance to ICBT, therein demonstrating a role for the IFNγ target gene product poly-ADP ribosyl polymerase 14 (PARP14). Although less studied than other PARPs, PARP14 has recently emerged as a promising therapeutic target in chronic inflammation. As a STAT6 transcriptional co-activator, PARP14 polarises immune responses towards those that are type 2 T helper (T_H_2) mediated^15,16^; while in IFNγ-treated macrophages, pro-inflammatory differentiation was suppressed by PARP14 through inhibition of STAT1 phosphorylation and down-regulation of STAT1 target genes^17^. Although PARP14 is an established oncoprotein^18,19^, the characterisation of its pro-or anti-inflammatory functions in the context of tumour immune evasion remains poorly characterised.

## Results

### Chronic IFNγ exposure drives resistance to α-PD-1 therapy and upregulates PARP14

Subcutaneous transplantation of mouse YUMM2.1 melanoma and CT26 and MC38 colon carcinoma cells into immunocompetent syngeneic mice gives rise to tumours that regress upon treatment with α-PD-1 antibodies^20,21^. However, chronic exposure of tumour cells to IFNγ limits the effectiveness of immune checkpoint inhibitors such as α-PD-1, driving resistance to treatment^11^. To validate the role of chronic IFNγ exposure in α-PD-1 therapy resistance, we implanted syngeneic mouse hosts subcutaneously with either IFNγ-naïve, bovine serum albumin (BSA)-exposed YUMM2.1, CT26 and MC38 cells or with the same cells pre-treated for 2 weeks with 50 IU/mL IFNγ. Once tumours were established, mice were subsequently treated with either α-PD-1 or IgG2a isotype control antibody (Fig. 1A). Chronic IFNγ pre-treatment did not affect tumour growth rates when mice were treated with control antibody (Extended Data Fig. 1A-C). Tumours derived from all three cell lines without chronic IFNγ pre-treatment demonstrated delayed growth in response to α-PD-1 therapy for at least one week post-treatment before mice eventually succumbed to progressive tumour growth. In contrast, IFNγ pre-treatment eliminated the ability of tumours to respond to α-PD-1 therapy significantly shortening survival (Fig. 1B-H), indicating that adaptation to IFNγ promotes α-PD-1 therapy resistance.

**Figure 1.**
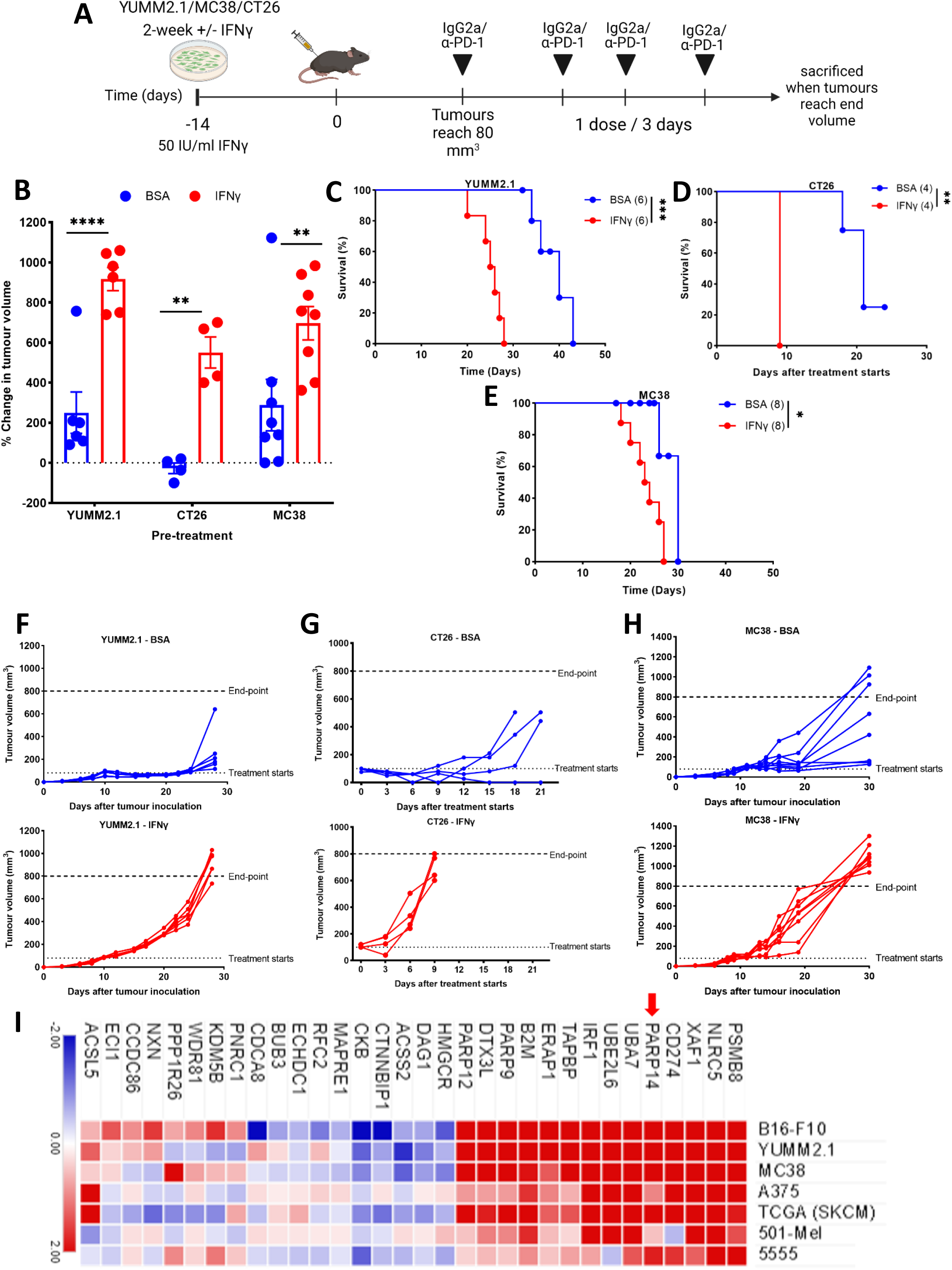
Chronic IFNγ exposure drives resistance to α-PD-1 therapy and upregulates PARP14. (A) YUMM2.1, CT26, and MC38 cells were implanted into 8–12-week-old wild-type syngeneic female mice after two-weeks pre-treatment with IFNγ (50 IU/mL) or BSA. Treatment with α-PD-1 antibody was initiated once tumour volume reached 80-100 mm^3^, with dosing every three days for a total of four doses. (B) The percentage change in tumour volume between the start of treatment and one week after the last α-PD-1 dose (YUMM2.1 and MC38) or the day that final dose was administered (CT26). (C-E) Kaplan-Meier survival plots of each animal receiving YUMM2.1 (C), CT26 (D), and MC38 (E) implants. (F-H) The growth curve of each YUMM2.1 (F), CT26 (G), and MC38 (H) tumour. Statistical significance determined by Unpaired t test or Log-rank (Mantel-Cox) test; *P < 0.05, **P < 0.01, ***P < 0.001. (I) Gene expression heatmap of differentially expressed genes (complete Euclidean HCL clustered; log2 fold change ≥ ±0.5; FDR ≤ 0.1) in mouse (B16-F10, YUMM2.1, MC38, 5555) or human (A375, 501-Mel,) tumour cell lines treated with chronic IFNγ (50 IU/mL for mouse and 20 IU/mL for human) compared to BSA treatment. Three independent cell line samples were sequenced for both conditions and the average for each cell line is shown. Heatmap also includes results from differential gene expression analysis of melanoma patient samples with the 15% highest IFNG expression levels relative to the 15% lowest (data retrieved from TCGA SKCM RNA sequencing data using Broad GDAC Firehose).

Chronic IFN exposure induces constitutive (ligand-independent) target gene expression through epigenetic reprogramming^13^. To identify mechanisms of IFNγ-driven adaptive resistance to α-PD-1, we exposed human (A375 and 501-mel) and mouse (B16-F10, MC38, 5555, and YUMM2.1) tumour cell lines to IFNγ (20 IU/mL for human and 50 IU/mL for mouse cell lines) or BSA continuously for 2 weeks, and subsequently performed RNA sequencing (RNA-seq). In addition to well-established IFNγ target genes such as *CD274, IRF1*, and *B2M*, three members of the PARP family, *PARP9, −12* and *−14*, were consistently upregulated in all cell models as well as in expression data from *IFNG*^high^ patient melanoma (comparing top 15% by *IFNG* expression to lowest 15% in the TCGA SKCM dataset) (Fig. 1I), suggesting that they may be direct IFNγ target genes. In agreement, we observed increased PARP14 levels in response to higher doses of IFNγ or repeat exposure in 6 human and mouse tumour cell lines (A375, 501-Mel, LOX-IMVI, CT26, MC38, and YUMM2.1) (Extended Data Fig. 1D, E). *In silico* analysis indicated the presence of multiple putative STAT1 binding sites on the PARP14 promoter (Extended Data Fig. 2A). Analysis of ChIP-seq data (retrieved from ENCODE project database)^22^ confirmed the binding of STAT1 near the transcription start site of PARP14 in IFNγ-treated cells (Extended Data Fig. 2B). In A375 cells transfected with a reporter plasmid in which *Gaussia* luciferase is regulated by the PARP14 promoter, luciferase activity increased with exposure to increasing IFNγ concentration. Furthermore, short interfering RNA (siRNA)-mediated STAT1 depletion in these cells impaired the ability of IFNγ to activate the reporter, confirming PARP14 induction through the IFNγ-STAT1 axis (Extended Data Fig. 2C). In keeping with PARP14 being an IFNγ target gene, the mRNA abundance of *PARP14* in both melanoma cells and patient samples (TCGA SKCM) positively correlated with that of *STAT1, IFNG*, and *CD274* (Extended Data Fig. 2D-G).

### PARP14 depletion or inhibition in tumour and host cells reverses adaptive resistance to α-PD-1 therapy

To address the role of PARP14 in chronic IFNγ-driven resistance to α-PD-1, we engineered YUMM2.1 and MC38 cells to express short-hairpin (sh) RNA targeting *PARP14* (shPARP14) for RNA interference-mediated down-regulation (Extended Data Fig. 3A). Tumour cells expressing shPARP14 showed significantly reduced PARP14 levels relative to cells expressing a non-target control shRNA (shNTC), even after a two-week IFNγ treatment (Extended Data Fig. 3A). We next implanted IFNγ pre-treated shNTC-or shPARP14-expressing YUMM2.1 and MC38 cells into mice and applied the same IgG2a or α-PD-1 treatment regimen described above (Fig. 2A). PARP14 depletion had no significant effect on the tumour formation or growth potential of YUMM2.1 or MC38 tumour cells in control IgG2a-treated mice (Extended Data Fig. 3B, C). However, PARP14 depletion restored responsiveness to α-PD-1 therapy (Fig. 2B-C and Extended Data Fig. 3D). Quantitation of *Parp14* mRNA expression in bulk tumour by RT-PCR analysis revealed that while expression was still significantly lower in endpoint YUMM2.1 tumours expressing shPARP14 compared to tumours expressing shNTC, this was not the case for MC38 tumours (Extended Data Fig. 3E), suggesting a selection for elevated *Parp14* expression in MC38 tumours treated with α-PD-1 and perhaps accounting for the less robust effect of PARP14 depletion in this model.

Despite only partially depleting PARP14 and in tumour cells alone, the above experiments indicated that PARP14 might mediate chronic IFNγ-induced adaptive resistance to α-PD-1 therapy. To explore this further and to demonstrate PARP14’s potential as a therapeutic target capable of modulating α-PD-1 sensitivity, rather than use shRNA to inhibit PARP14, we treated mice implanted with IFNγ pre-treated YUMM2.1, CT26, or MC38 cells with α-PD-1 in combination with RBN012759, a highly selective PARP14 catalytic inhibitor (PARP14i) (Fig. 2D). According to previous findings, twice daily dosing of mice with 500 mg/kg RBN012759 achieves stable PARP14 suppression without adverse effects^23,24^. Pharmacological inhibition of PARP14 strongly synergised with α-PD-1, with tumour regression and significantly extended survival observed in all three models (Fig. 2E, F and Extended Data Fig. 4A-E). 25% of mice bearing YUMM2.1 tumours treated with a combination of α-PD-1 and PARP14i exhibited durable tumour regression (up to 60 days post-treatment). Additionally, at 2 months post cessation of combination therapy, all long-term survivors rejected the re-implantation of chronic IFNγ pre-treated YUMM2.1 cells (Extended Data Fig. 4F), indicating the induction of anti-tumour immune memory. Next, we addressed the extent to which CD8+ T cells control tumour growth in α-PD-1/PARP14i combination-treated animals (Extended Data Fig. 4G). We found that depleting CD8+ cells through systemic administration of α-CD8 antibody permitted progression of combination therapy-treated tumours (Extended Data Fig. 4H-J).

**Figure 2.**
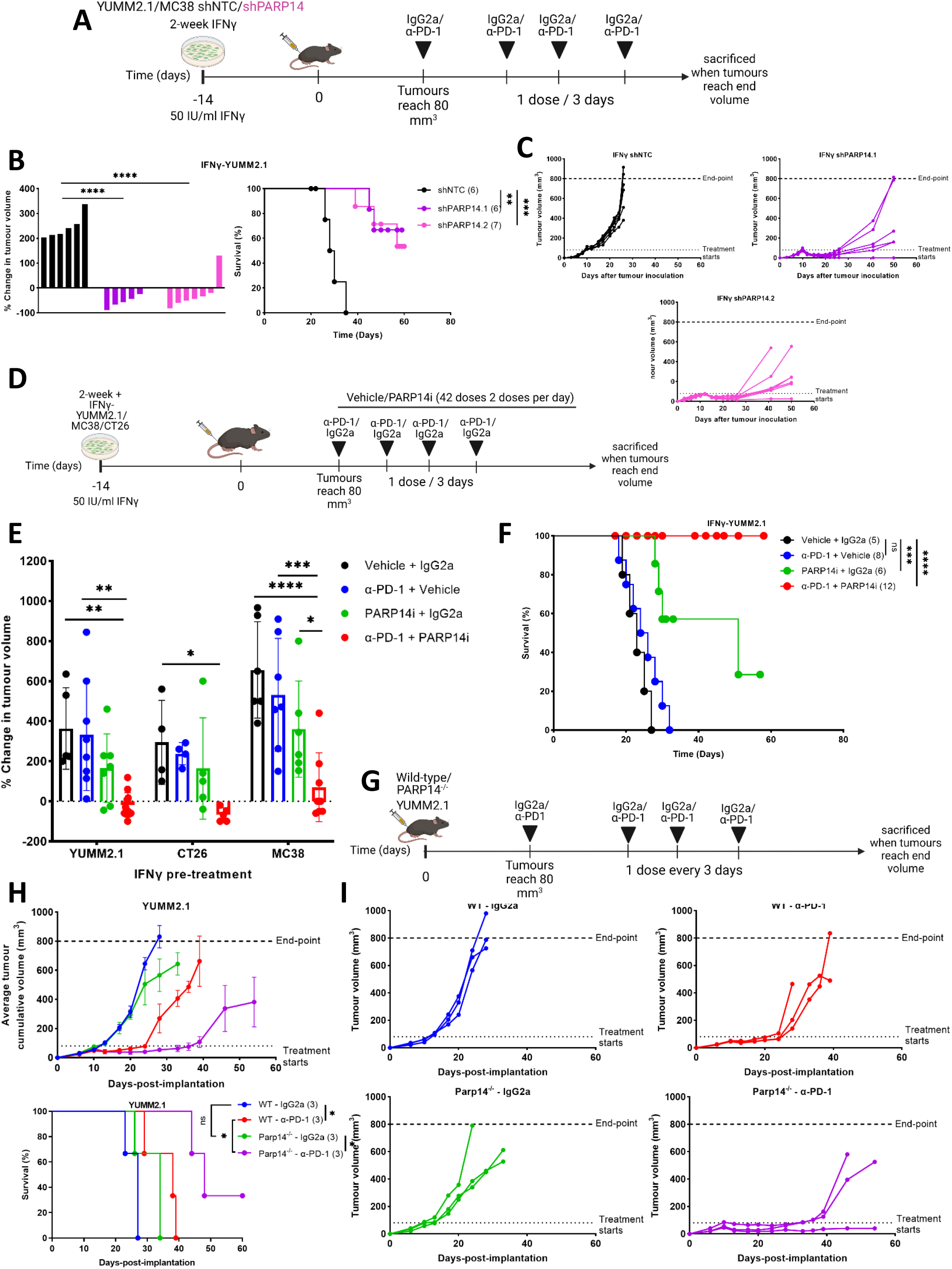
PARP14 depletion in tumour cells or pharmacological antagonism reverse adaptive resistance to α-PD-1 therapy while lack of PARP14 in host cells augments efficacy. (A) Chronic IFNγ pre-treated YUMM2.1 cells expressing two independent PARP14-targeting shRNAs (shPARP14) or a non-target control shRNA (shNTC) were subcutaneously implanted into 8–12 weeks-old wild-type C57BL/6 female mice. Treatment of tumour-bearing mice was initiated once tumour volume reached ~80 mm^3^, with dosing every three days for a total of four doses. (B) The percentage tumour volume change between the first α-PD-1 treatment dose and the final dose (left) and Kalan-Meier plots for these mice (right). Kaplan-Meier plots statistical significance determined by Log-rank (Mantel-Cox) test; **P < 0.01, ***P < 0.001. (C) Tumour growth rate for each treatment group. (D) Chronic IFNγ pre-treated YUMM2.1/MC38/CT26 cells were subcutaneously implanted into 8–12-week-old wild-type syngeneic female mice. Treatment with either α-PD-1 or IgG2a antibody was initiated once tumour volume reached 80-100 mm^3^, with antibodies administered every three days for a total of four doses. In parallel, the animals also received two daily doses of the PARP14 inhibitor (PARP14i) RBN012759 or vehicle for a total of three weeks. (E) The percentage change in tumour volume between the first dose of treatment and the administration of the final α-PD-1 dose of mice receiving implants of chronic IFNγ pre-treated YUMM2.1, CT26, and MC38. Statistical significance determined by one-way ANOVA; *P < 0.05, **P < 0.01, ****P < 0.0001. (F) Kaplan-Meier plots of mice receiving implants of chronic IFNγ pre-treated YUMM2.1. Statistical significance determined by Log-rank (Mantel-Cox) test; NS P > 0.05, ***P < 0.001, ****P < 0.0001. (G) IFNγ-naïve YUMM2.1 cells were subcutaneously implanted into 8–12-week-old wild-type or Parp14^−/−^ C57BL/6 female mice, followed by antibody treatment as described above. (H) Average cumulative tumour volume growth curve (top) and Kaplan-Meier plots (bottom). Kaplan-Meier plots statistical significance determined by Log-rank (Mantel-Cox) test; NS P > 0.05, *P < 0.05. (I) Tumour growth rate for each treatment group.

PARP14 expression is not restricted to tumour cells; it is also expressed in immune cells and multiple other normal cell types^15,16,25^. Previous findings also demonstrated that PARP14 might perform cancer-promoting functions in stromal cells present in the TME^16,17,23^. To determine whether endogenous PARP14 expression by host cells could affect the overall efficacy of α-PD-1 therapy, *Parp14*-KO or wild-type mice were subcutaneously injected with YUMM2.1 cells and subsequently treated with four doses of α-PD-1 (Fig. 2G). We observed significantly improved tumour growth suppression and overall survival was significantly enhanced in the knockout mice compared to wild-type mice (Fig. 2H, I), indicating that PARP14 loss in host cells was also beneficial to α-PD-1 response.

### Chronic IFNγ exposure reshapes the tumour immune infiltrate through PARP14

To assess whether and how chronic IFNγ signalling affects tumour immune cell infiltration and the contribution of PARP14 to this, we profiled the immune infiltrate of subcutaneous tumours derived from IFNγ-naïve YUMM2.1 cells expressing shNTC or chronic IFNγ pre-treated YUMM2.1 cells expressing either shNTC or shPARP14 (Fig. 3A) by flow cytometry using fluorescent labelling for a panel of T cell markers including TCRαβ, TCRγδ, CD45, CD25, and FoxP3 (Extended Data Fig. 5 for gating strategy). Tumours derived from chronic IFNγ-pre-treated YUMM2.1 cells exhibited a significantly lower percentage of T cells (TCRαβ^+^) and a higher percentage of regulatory T (T_reg_) cells (Foxp3^+^, CD25^high^) relative to tumours derived from either IFNγ-naïve cells or shPARP14-expressing IFNγ-pre-treated YUMM2.1 cells (Fig. 3B), implying that PARP14 might contribute to the immunosuppressive tumour microenvironment induced by chronic IFNγ pre-treatment. Next, we assessed whether PARP14i alone or in combination with α-PD-1 could reverse the immunosuppressive effects of chronic IFNγ pre-treatment (Fig. 3C). Compared to control-treated tumours, the combination of α-PD-1 and PARP14i elicited the greatest increase in the number of CD45^+^ immune cells relative to tumour mass, with an increased percentage of CD8^+^ T cells and a decreased percentage of T_reg_ cells, leading to a significant increase in the ratio of CD8^+^ Granzyme B^+^ (GzmB^+^) cytotoxic lymphocytes (CTLs) to T_reg_ cells (Fig. 3D and Extended Data Fig. 6 for gating strategy).

**Figure 3.**
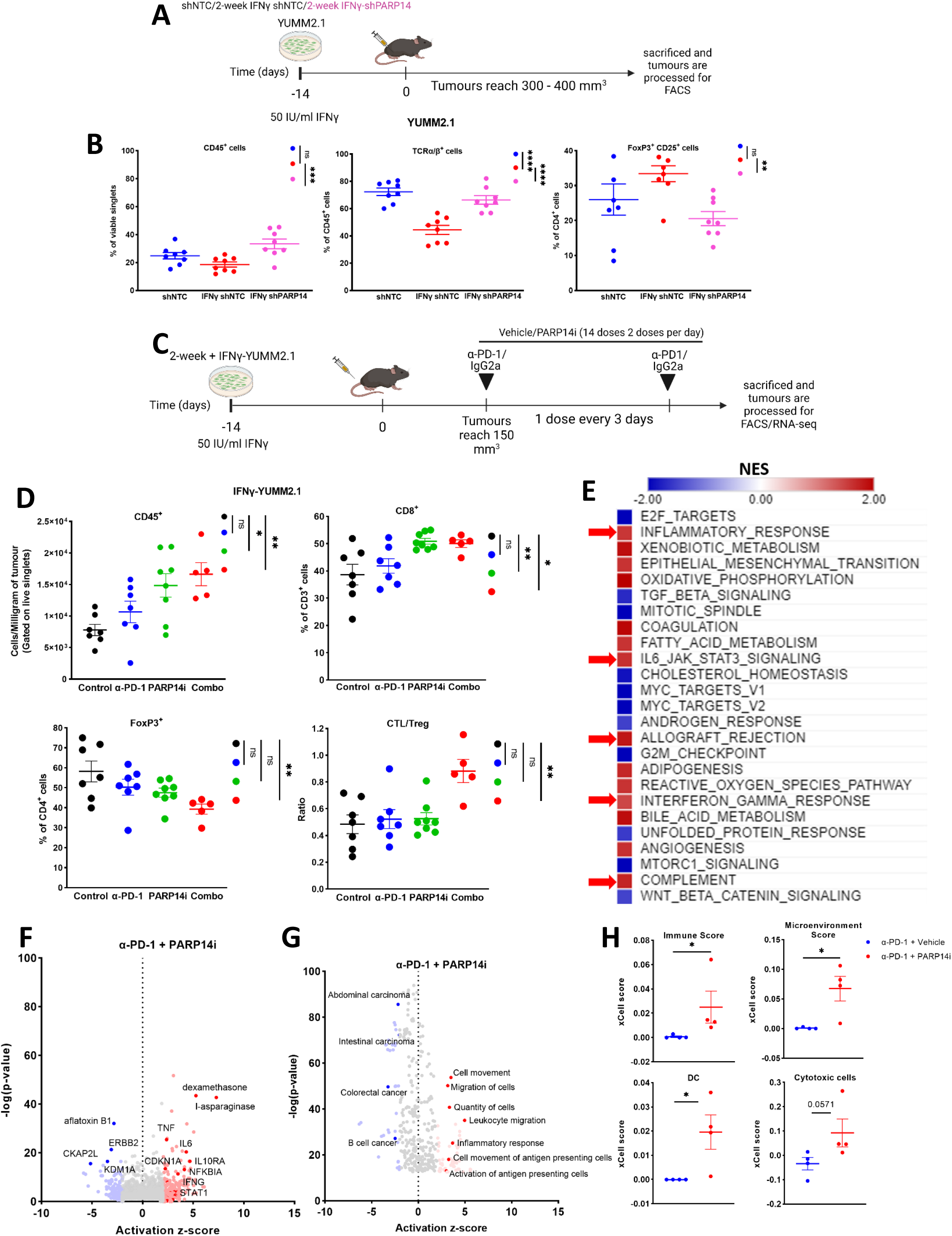
PARP14 depletion or inhibition reverses chronic IFNγ driven immune regulatory effects. (A) 8–12-week-old wild-type C57BL/6 mice were subcutaneously implanted with IFNγ-naïve YUMM2.1 cells expressing shNTC or chronic IFNγ pre-treated YUMM2.1 cells expressing shNTC or shPARP14. Tumours were allowed to grow to 300-400 mm^3^ and then dissected and disaggregated for analysis by flow cytometry. (B) Populations of total immune cells (CD45^+^), T cells (TCRαβ^+^), and regulatory T cells (T_reg_ cells; CD25^+^FoxP3^+^) in the tumour infiltrate. (C) 8–12-week-old wild-type C57BL/6 mice were subcutaneously implanted with chronic IFNγ pre-treated YUMM2.1 cells. Treatment with either α-PD-1 or IgG2a antibody was initiated once tumour volume reached 150 mm^3^ (two doses, three days apart) and PARP14i or vehicle (two doses daily for a week). At the end of the treatment period, tumours were dissected and disaggregated for analysis by flow cytometry and RNA-seq. (D) Populations of total immune cells (CD45^+^), cytotoxic T cells (CD8^+^), Treg cells (FoxP3^+^), and the ratio of CD8^+^GzmB^+^ cytotoxic lymphocytes (CTLs) to T_reg_ cells in the tumour infiltrate. Data are presented as mean ± S.E.M. for ≥ 6 independent samples. Statistical significance determined by one-way ANOVA; NS P > 0.05, *P < 0.05, **P < 0.01, ***P < 0.001, ****P < 0.0001. (E) Gene set enrichment analysis (GSEA) hallmark results for targets upregulated (red) or downregulated (blue) by α-PD-1/PARP14i combination treatment. Normalised enrichment score (NES) ≥ ±0.5; FDR ≤ 0.25. Red arrows indicate hallmark gene sets of inflammatory response, IL6-JAK-STAT3 signalling, allograft rejection, interferon-gamma response, and complement. (F, G) Ingenuity Pathway Analysis (IPA) was applied to identify up-or down-regulation of upstream regulators and disease-related or functional pathways in tumours receiving α-PD-1/PARP14i combination treatment. Control was α-PD-1 monotherapy. Results are displayed with their P-value (-log(P-value)) and activation z-score. (H) RNA-seq results were analysed by cell type enrichment analysis (xCell score), with scores shown for immune, microenvironment, dendritic cell (DC), and cytotoxic T cells. Statistical significance determined by Unpaired two-tailed Mann-Whitney test; NS P > 0.05, *P < 0.05, **P < 0.01, ***P < 0.001, ****P < 0.0001.

We also investigated gene expression differences in tumours derived from chronic IFNγ-pre-treated YUMM2.1 cells treated with α-PD-1 monotherapy versus α-PD-1 + PARP14i combination therapy by sequencing mRNA from bulk tumours. Gene set enrichment analysis (GSEA) revealed that the combination therapy upregulated numerous inflammatory signalling pathways (Fig. 3E). Furthermore, Ingenuity Pathway Analysis indicated that *STAT1, IFNG*, and *TNF* responses were strongly activated when PARP14 was also inhibited (Fig. 3F). Additionally, leukocyte migration and activation of antigen-presenting cells were strongly activated and tumourigenesis-related processes strongly down-regulated following PARP14i treatment (Fig. 3G). We also performed computational immunophenotyping using xCell cell-type enrichment analysis^26^, which indicated that immune score, microenvironment score, dendritic cells (DC), and cytotoxic cells were upregulated in tumours undergoing combination therapy (Fig. 3H), consistent with PARP14i contributing to increased immune infiltration. Collectively, these data show that PARP14 antagonism potentiates the immunostimulatory effect of α-PD-1 in an otherwise immunosuppressive tumour microenvironment established by chronic IFNγ-signalling.

### PARP14 is a negative feedback regulator of IFNγ signalling

PARP14 down-regulates STAT1 in IFNγ-stimulated macrophages and thereby antagonises IFNγ-induced macrophage polarisation^17^. We hypothesised that PARP14 might also act as a feedback inhibitor in tumour cells, thereby antagonising IFNγ-stimulated tumour cell immunogenicity. In keeping with this hypothesis, we found that phospho-STAT1 (pSTAT1), STAT1, and STAT1 target gene products PD-L1, MHCI, TAP1, and TAP2 were enriched in shPARP14-expressing, chronic IFNγ-treated YUMM2.1 and MC38 cells compared to shNTC-expressing cells (Extended Data Fig. 3A). Similarly, pharmacological antagonism of PARP14 using either RBN012759 or the proteolysis targeting chimera (PROTAC) PARP14 inhibitor RBN012811^23,27^ at nanomolar concentrations in chronic IFNγ-treated MC38, YUMM2.1, 501-Mel, or A375 cells resulted in elevated levels of pSTAT1 and STAT1 target gene products without perturbing the growth of these cell lines (Fig. 4A-C). Intriguingly, while the expected depletion of PARP14 protein occurred following the degradation-inducing RBN012811 treatment, application of the catalytic inhibitor RBN012759 led to elevated levels of PARP14 protein, consistent with PARP14 being itself a STAT1 target. Moreover, RNA-seq and subsequent GSEA revealed that PARP14 inhibition enhanced inflammatory signalling (Fig. 4D). Quantitative PCR confirmed a significant increase of mRNA expression for the chemokine ligands *Cxcl10* and *Cxcl11* (Fig. 4E), supporting our hypothesis that PARP14 inhibition enhances IFNγ signalling in tumour cells and upregulates immune cell infiltration into tumours.

**Figure 4.**
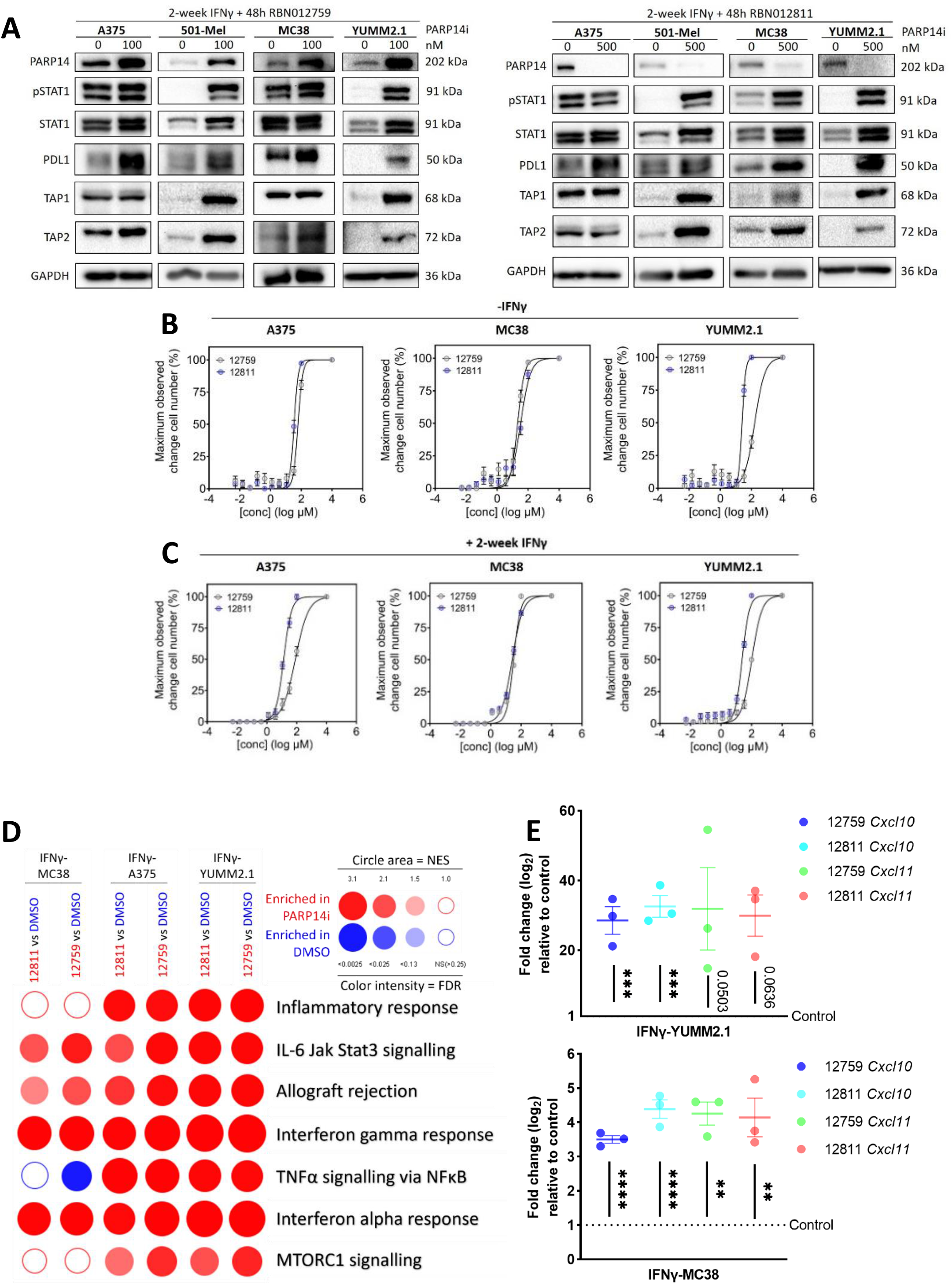
PARP14 is a negative feedback regulator of IFNγ signalling. (A) Chronic IFNγ pre-treated A375, 501-Mel, MC38, or YUMM2.1 cells were treated for 48 hours with PARP14 pharmacological inhibitors RBN012759 (left) and RBN012811 (right). Expression of PARP14, pSTAT1, STAT1 and STAT1 target proteins is shown by western blot, with GAPDH applied as a loading reference. (B) Graphs showing the relative maximum observed change in cell number as determined by crystal violet following 48-hour treatment with varying concentrations of PARP14 inhibitors (mean, n>3). (D) BubbleGUM GSEA based on RNA-seq data, depicting hallmark processes enriched in chronic IFNγ pre-treated tumours treated with RBN012759 or RBN012811 versus control (DMSO). The circle area depicts the NES, and colour intensity depicts the FDR, with ≤ 0.25 classed as significant. (E) RT-qPCR analysis of *Cxcl10* and *Cxcl11* mRNA expression levels in YUMM2.1 and MC38 cells. Data show mean ± S.E.M. for three independent samples. Statistical significance determined by unpaired t-test; NS P>0.05, **P<0.01, ***P<0.001, ****P < 0.0001.

### PARP14 levels are augmented in tumours spontaneously relapsing after α-PD-1 treatment wherein it mediates resistance

To address the role of PARP14 in spontaneously arising adaptive resistance to α-PD-1 therapy, we firstly validated whether PARP14 expression could be induced by α-PD-1 therapy. Following establishment of tumours derived from IFNγ-naïve YUMM2.1 and MC38 cells, mice were treated with three doses (spaced three days apart) of IgG2a or α-PD-1 antibodies. Tumours were harvested within 24 hours of the last dose of treatment. Quantitative PCR did not reveal a significant upregulation of *Parp14* in bulk-tumour mRNA from α-PD-1 on-treatment tumours compared to control tumours (Extended Data Fig. 7A). However, when we grouped YUMM2.1 and MC38 tumour specimens by the level of *Ifng* mRNA (*Ifng*^high^ versus *Ifng*^low^) regardless of treatment, *Parp14* expression was significantly higher in *Ifng*^high^ tumours. We made similar observations for *Stat1* and other STAT1 target genes including *Irf1, Cxcl10*, and *Cxcl11* (Extended Data Fig. 7B), which suggested that PARP14 was induced specifically in IFNγ-inflamed tumours but independently of α-PD-1. Turning to expression data from human melanoma biopsies^28^, we observed a modest but significant increase in *PARP14* mRNA in α-PD-1 on-treatment melanoma biopsies compared to pre-treatment biopsies that correlated with increased *IFNG* and *STAT1* mRNA (Extended Data Fig. 7C).

Despite PARP14 induction in tumours derived from IFNγ-naïve YUMM2.1 cells not depending on α-PD-1, we found that *Ifng, Stat1* and their target genes, including *Parp14*, were significantly increased in bulk tumour mRNA of tumours relapsing following α-PD-1 treatment compared to control tumours (Fig. 5A). We, therefore, addressed whether tumours that relapsed following α-PD-1 treatment were sensitive to PARP14 inhibition. IFNγ-naïve YUMM2.1 cells were implanted subcutaneously, and subsequently tumour-bearing mice received two doses of IgG2a or α-PD-1 antibodies three days apart (Fig. 5B). Following the final dose of α-PD-1, tumours were permitted to regrow to their pre-treatment size before further treatment commenced. At this point, we initiated one week of treatment with either vehicle, a further two doses of α-PD-1 or PARP14i (14 doses). We observed that 33% of these relapsing mice experienced complete regression within one week of initiating PARP14i treatment, with an overall significant decline in average tumour cumulative volume (Fig. 5C). In contrast, continuing treatment with α-PD-1 antibodies had no further effect on tumour growth (Extended Data Fig. 7D). We also noted that tumours derived from IFNγ-naïve YUMM2.1 cells were largely insensitive to PARP14i alone treatment (Extended Data Fig. 7E). The growth of tumours derived from IFNγ-naïve CT26 cells that regrew following α-PD-1 administration was also suppressed by PARP14i treatment (Extended Data Fig. 7F).

**Figure 5.**
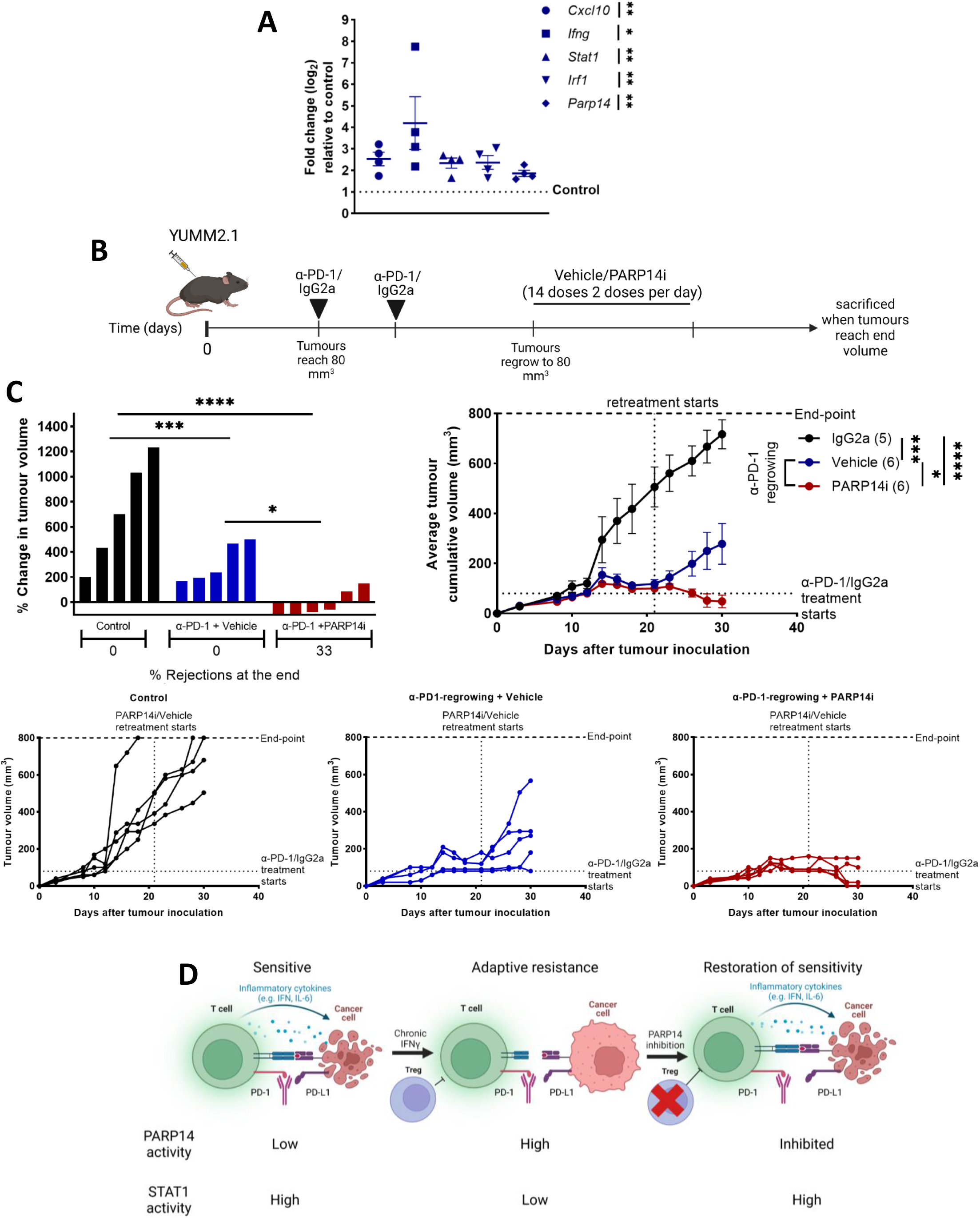
PARP14 levels are augmented and mediate resistance in tumours spontaneously relapsing after α-PD-1 treatment. (A) RT-qPCR analysis of *Cxcl10, Ifng, Stat1, Irf1*, and *Parp14* mRNA expression levels in YUMM2.1 tumours which regrew after α-PD-1 antibody treatment compared to IgG2a-treated tumours (control). Statistical significance determined by unpaired t-test; *P < 0.05, **P < 0.01. (B) 8–12-week-old wild-type C57BL/6 mice were subcutaneously implanted with YUMM2.1 cells. α-PD-1 antibody treatment (two doses, three days apart) was initiated once tumour volume reached ~80 mm^3^. Following subsequent tumour regrowth to ~80 mm^3^, mice were treated with two daily doses of PARP14i or vehicle for a week. (C) The percentage change in tumour volume between the start of α-PD-1 treatment and day 30 post-tumour-implantation (top left), average tumour cumulative volume growth curve (top right), and growth curves for each tumour (bottom). Statistical significance determined by one-way ANOVA; *P < 0.05, ***P < 0.001, ****P < 0. 0001. (D) Proposed model of response to PD-1 immune checkpoint inhibition restoration by PARP14 inhibition in tumours with resistance driven by IFNγ.

## Discussion

IFNγ is a potent stimulant of cellular immunity, inducing the expression of several gene products involved in antigen processing and presentation, chemotaxis, inflammation, and antibacterial and antiviral immune responses^29^. However, the role of IFNγ in ICBT resistance remains controversial, mirroring the complexity of IFNγ signalling in tumorigenesis and immune evasion^29^. In this study, we aimed to identify novel mechanisms of IFNγ-driven immune evasion and ICBT resistance and identified the IFNγ target gene *PARP14* as a critical mediator (summarised in Figure 5D).

RNA-seq analysis showed upregulation of *PARP14* in multiple human and mouse tumour cell lines chronically exposed to IFNγ as well as in *IFNG*^high^ melanoma tumours, indicating that *PARP14* is an IFNγ target gene. Indeed, our data imply that *PARP14* is a direct STAT1 target. Moreover, PARP14 appears to negatively regulate IFNγ signalling and responses, consistent with PARP14 being a negative feedback regulator of IFNγ signalling. In addition, PARP14 interacts with proteins that are commonly co-expressed following IFN treatment and also interact with STAT1, thereby potentially regulating the IFN-induced interactome^30^. PARP9 is such a protein, which promotes STAT1 activation and pro-inflammatory gene expression in IFNγ-treated macrophages^17^ as well as in pancreatic epithelial and cancer cells^31^. In contrast, Iwata and colleagues suggested that PARP14 suppresses STAT1 by ADP-ribosylating STAT1 to prevent its phosphorylation in response to IFNγ^17^. In keeping with PARP14 being a negative feedback regulator of IFNγ signalling, we found that PARP14 silencing or pharmacological antagonism in tumour cells enhanced STAT1 phosphorylation and expression of STAT1 target gene products, including PARP14 itself. Further, RNA pol II ChIP-seq employed by Riley and colleagues identified 2,744 genes down-regulated by PARP14, including genes encoding components of the antigen processing and presentation machinery^15^.

For our *in vivo* experiments, we employed YUMM2.1, CT26 and MC38 syngeneic mouse tumour models, which initially respond to α-PD-1 therapy but subsequently relapse, MC38 more quickly than YUMM2.1 or CT26. Chronic IFNγ treatment before implantation rendered tumours resistant from the outset to α-PD-1 therapy, but response to α-PD-1 could be restored by depleting PARP14 in tumour cells or pharmacologically antagonising PARP14 systemically. Notably, combining PARP14i with α-PD-1 treatment potently suppressed tumour growth, with occasional complete tumour regression, dependent on CD8^+^ cells. As healthy tissues also express PARP14, we also treated *Parp14*^−/−^ mice with α-PD-1 therapy and found that the absence of PARP14 in host cells also improved the control of YUMM2.1 tumour growth. Cho and colleagues found that while *Parp14*^−/−^ mice exhibited normal total cell numbers in the thymus, spleen, and lymph nodes, they had increased proportions of CD8^+^ T cells^18^. This might explain the moderately better response of these mice to α-PD-1 therapy. However, the impact of PARP14 expression in host cells—especially in immune cells—on tumour growth and α-PD-1 therapy efficacy warrants further investigation.

We also explored the effects of IFNγ preconditioning and PARP14 depletion on the composition and activation status of the tumour immune infiltrate. We found that chronic IFNγ treatment of YUMM2.1 cells drastically remodelled the TME. Specifically, these tumours were infiltrated by significantly fewer CD45^+^ immune cells, including TCRαβ^+^ cells, whereas T_reg_ cells were proportionately enriched, mirroring the ability of lymph node metastases that are likewise exposed to chronic IFNγ stimulation to induce or promote the recruitment of T_reg_ cells^32^. Importantly, PARP14 silencing in IFNγ pre-treated tumours reversed these alterations in the TME. These effects of silencing PARP14 on the TME were consistent with restoring sensitivity to α-PD-1, as pre-existing immune infiltration and high expression levels of immune-related genes and low numbers of T_reg_ cells predict good response to α-PD-1 therapy in patients^33–36^. Subsequently, we determined that the synergy between α-PD-1 therapy and PARP14i may also be explained by effects on the composition of the tumour immune infiltrate. Using the IFNγ pre-treated YUMM2.1 model, we found that the combination therapy led to an increase in intratumoral CD45^+^ immune cells and CD8^+^ T cells and a decreased frequency of T_reg_ cells. While this drug combination did not appear to affect T cell exhaustion (data not shown), potential effects on stemness warrant further investigation. Our study focused on the ability of our therapeutics to influence cytotoxic T-cell responses; however, mounting evidence suggests that other immune cells are also important for α-PD-1 efficacy. For example, NK cells are essential for response to PD-1/PD-L1 blockade in some models^37^. Moreover, PARP14 facilitates polarisation of macrophages into an M2-like state and of CD4^+^ T cells to the T_H_2 lineage, both of which are cancer-promoting immune cell subtypes^16,17^. The effect of chronic IFNγ exposure on and possible contribution of PARP14 to tumour infiltration by myeloid and helper T cell populations also merits further investigation.

The contribution of IFNγ signalling to ICBT resistance remains controversial. The induction of IFNγ and its target genes are sensitive and robust prognostic markers of ICBT response^8,37^. Contrasting *IFNG* and *STAT1* mRNA abundance in pre-versus on-treatment melanoma biopsies^28^ also supported that IFNγ-signalling is induced by ICBT. As such, insensitivity to IFNγ might be predicted to benefit tumour cells in the context of ICBT. In keeping with this, Gao and colleagues showed that tumours from patients resistant to α-CTLA-4 therapy harboured genomic defects in IFNγ pathway components, including copy-number loss of IFNγ pathway genes (e.g., *IFNGR1/2, IRF1*, and *JAK2*) and amplification of IFNγ pathway inhibitors (e.g., *SOCS1* and *PIAS4*)^9^, while whole-exome sequencing of patient biopsies revealed loss-of-function mutations in JAK1 and JAK2 in patients with primary and acquired resistance to PD-1 blockade^10,38^. Although these studies suggest that inactivation of IFNγ Pathway components is a major cause of ICBT resistance, this conclusion was based on limited patient samples, and subsequent studies failed to detect these changes at a significant frequency in larger populations^28,39,40^. Furthermore, all the cell lines used in our study respond robustly to IFNγ, suggesting that IFN signalling is preserved despite immunoediting. Further, more abundant IFNγ target gene products in plasma from patients undergoing ICBT predicted relapse and poorer survival^12^. Negative feedback inhibition by downstream IFNγ pathway components such as PARP14 may be the more common mechanism of adaptation to chronic IFNγ signalling induced by ICBT than inactivating mutations in upstream components of the pathway.

Unsurprisingly for an IFNγ target gene, induction of *PARP14* mRNA mirrored *IFNG* and *STAT1* mRNA in on-treatment melanoma biopsies. However, *PARP14* mRNA levels could not predict the depth or duration of clinical response among responders to ICBT in the Riaz *et al*. cohort^28^ (data not shown). This may reflect several confounding factors, including a limited number of samples available for analysis; highly heterogeneous tumours in which only small regions were sampled; sampling of different lesions pre- and on-treatment; and finally, biopsies that were not necessarily sampled at the point of relapse, which is when our *in vivo* data indicate that *PARP14* induction may be at its highest and exerting its greatest influence. Therefore, a more robust longitudinal examination of *PARP14* induction in patient biopsies undergoing α-PD-1 therapy is needed, although this may be difficult to reconcile with patient care.

Although PD-L1 induction is the most well-established mechanism by which IFNγ signalling restores immune homeostasis and drives tumour immune evasion^41–43^, multiple PD-L1-independent immunomodulatory mechanisms also exist^44–46^. Depletion of the JAK-STAT signalling regulator LNK impaired tumour growth and potentiated α-PD-1 responses by relieving LNK-mediated STAT1 inhibition^42^. Similarly, inhibition of RIPK1 enhances STAT1 signalling and the activation of cytotoxic T cells contributing to anti-tumour activity^47^. Chronic IFNγ signalling consistently induced expression of Qa-1b/HLA-E, a ligand for the cytotoxic lymphocyte inhibitory receptor NKG2a/CD94, in mouse tumour models, thereby conferring resistance to α-PD-1 therapy^12^. The application of PD-L1-targeting therapies in combination with treatments that act independently of PD-L1 is, therefore, a promising strategy to overcome ICBT resistance.

The combination of PARP inhibitors with ICBT has recently emerged as a promising therapeutic strategy for cancer, with PARP1/2/3 inhibitors, such as the FDA-approved drugs niraparib, olaparib, and rucaparib now at the forefront of clinical investigations^48,49^. It is becoming increasingly evident that the anti-cancer effects of PARP inhibitors go beyond their direct cytotoxic effects and that these drugs may also enhance α-PD-1 efficacy by activating the stimulator of interferon genes (STING) independently of BRCA status^50^. This is achieved through the generation of cytosolic double-stranded DNA fragments, which bind cyclic GMP-AMP synthase (cGAS) and activate STING, thereby inducing a type I IFNα/β response. Consequently, chemokine secretion and subsequent T cell infiltration are enhanced^50^. Notably, the PARP1 inhibitor talazoparib (BMN 673) elicited increased numbers of peritoneal CD8^+^ T cells and NK cells in an ovarian cancer mouse model and increased infiltration into *ex vivo* spheroids, in addition to increasing IFNγ and TNF-α production levels^51,52^. The finding that PARP inhibitors upregulate PD-L1 in cancer cells further supports the rationale of combining PARP inhibition with α-PD-1 therapy for the treatment of ovarian, breast, and non-small cell lung cancer^53^.

In addition to these established FDA-approved treatments, targeting other PARP family members can also enhance anti-cancer immune responses. The demonstration of both cancer cell-autonomous effects and anti-tumour immunity induced by enhanced IFN signalling upon application of our highly selective PARP7 inhibitor (RBN-2397)^54^, lead to a phase 1 clinical study (NCT04053673) of this drug for patients with advanced solid tumours. Indeed, Falchook and colleagues showed upregulated *CXCL10* mRNA levels in tumours of patients treated with RBN-2397, accompanied by an increase in CD8^+^ GzmB^+^ T cells^55^. Unlike PARP1 and PARP7, Our data suggests that PARP14 inhibitor treatment would not be efficacious as a monotherapy, as PARP14 knockdown or antagonism did not affect the ex vivo growth of the tumour cell lines we evaluated, nor did monotherapy significantly alter their tumour growth potential, even following chronic IFNγ stimulation. Thus, despite mediating immune evasion driven by chronic IFNγ-signalling, PARP14 activity is redundant to other factors, notably PD-1 signalling.

In conclusion, we identified PARP14 as a key mediator of IFNγ-driven ICBT resistance. We were able to demonstrate that down-regulation or inhibition of PARP14 could promote IFNγ-STAT1 signalling, convert an immunosuppressive TME into a more immunostimulatory state, and reinstate sensitivity to α-PD-1, providing a strong rationale for combining PARP14-targeting interventions with α-PD-1 therapy.

## Methods

### Mouse tumour implant study design

Mice were housed in the Biological Services Facility of The University of Manchester on a 12/12-hour light/dark cycle, and given unlimited access to food (Bekay, B&K Universal, Hull, UK) and water. All procedures were approved by the institution’s review board and performed under relevant Home Office licenses according to the UK Animals (Scientific Procedures) Act, 1986. Female, 8–12-week-old C57BL/6 or BALB/c mice were purchased from ENVIGO and allowed at least 1 week to acclimatise; *Parp14*^−/−^ mice were originally provided by Dorian Haskard (Imperial College) and subsequently bred in-house. YUMM2.1 cells (7 × 10^6^ cells), CT26 (1 × 10^6^ cells), and MC38 cells (3 × 10^5^ cells) in 100 µL serum-free RPMI-1640 were subcutaneously injected into the left flank of mice under isoflurane anaesthesia. Tumour size (calculated by multiplication of height, width, and length calliper measurements) and mouse weight were monitored three times per week (every 2-3 days). When tumours reached an average volume of 80-100 mm^3^, mice were administered with up to four doses of 300 µg of α-PD-1 antibody (BioXCell) or rat isotype control antibody IgG2a (BioXCell) in 100 µL InVivoPure pH 7.0 Dilution Buffer (BioXCell) via intraperitoneal (i.p.) injection administered at 3–4-day intervals. Mice were also administered vehicle or 500 mg/kg of RBN012759 by oral gavage twice a day (BID). RBN012759 was dissolved in 0.5% methylcellulose (Sigma-Aldrich) + 0.2% Tween 80 (Sigma-Aldrich). Each dose was delivered in a volume of 0.2 mL/20 g mouse (10 mL/kg) and adjusted for the last recorded weight of individual animals. Mice were monitored and body weight was measured daily. Mice were culled once tumours reached 800 mm^3^, our pre-determined experimental endpoint aligning with the principles of the 3Rs (Replacement, Reduction and Refinement) for improving animal welfare. A sample size of n ≥ 3 per group was used throughout to achieve a statistical significance of *P* < 0.05. Mice were randomised into treatment groups. All tumours were included for analysis. Differences in survival were determined using the Kaplan-Meier method, and the P-value was calculated by the log-rank (Mantel-Cox) test using GraphPad Prism version 9.0c. A mixed-effect linear model using GraphPad Prism (version 9.0) was used to determine differences in growth curves. For the tumour response plots, the percentage of tumour volume change between a specific date of post-treatment and the start of the treatment was plotted for each tumour. The significance of all one-way comparisons was determined using one-way ANOVA. For non-parametric data, the unpaired t-test or Mann-Whitney test was used.

### Reagents

Recombinant human IFNγ (PHC4031) and recombinant mouse IFNγ (PMC4031) were purchased from Gibco and used at the indicated concentrations. The PARP14 inhibitors RBN012759^23^ and RBN012811^27^ were kindly provided by Ribon Therapeutics and used at the indicated concentrations.

### Cell lines

The human melanoma cell lines A375, LOX-IMVI and 501-Mel (provided by Claudia Wellbrock, The University of Manchester), Lenti-X 293T cells (provided by Angeliki Malliri, The University of Manchester), 5555, B16-F10, and MC38 cells (provided by Santiago Zelenay, The University of Manchester) and YUMM2.1 cells (provided by Richard Marais, The University of Manchester) were maintained in RPMI-1640 (Sigma-Aldrich) supplemented with 10% foetal bovine serum (FBS; Life Technologies) and 1% penicillin-streptomycin (P/S; Sigma-Aldrich). All cells were maintained under standard conditions at 37°C in a 5% CO_2_ humidified incubator and passaged before reaching confluency. Cell cultures were routinely tested for mycoplasma contamination by PCR.

### Cell proliferation

Cells were fixed and stained with 0.5 %w/v crystal violet (Sigma) in 4% paraformaldehyde (PFA) in phosphate-buffered saline (PBS) for at least 30 minutes. Fixed cells were solubilised in 2% sodium dodecyl sulphate (SDS) in PBS and absorbance was measured at 595 nm using Biotek Synergy™ H1 Hybrid Multi-Mode Reader.

### Gene silencing

For siRNA-mediated silencing of STAT1, cells were seeded in 6-well plates (5 × 10^5^ cells/well) and incubated overnight. The next day, cells were transfected with siRNAs using lipofectamine RNAiMAX (Life Technologies) transfection reagent according to the manufacturer’s guidelines. After eight hours of incubation with the transfection mixture, the cell culture medium was replaced, and cells were incubated for 1-3 days at 37°C. For shRNA-mediated silencing of PARP14, Lenti-X 293T cells were seeded in T75 flasks (5 × 10^6^ cells/flask) and incubated overnight. The next day, cells were transfected with 4.5 μg of the respective shRNA/overexpressing vector, 6 μg of psPAX2 (12260; Addgene), and 3 μg of pVSVg (8454; Addgene); FuGENE HD transfection reagent (Promega) was used for the transfections. All shRNAs were cloned in pLV-EGFP lentiviral transfer vectors (VectorBuilder); the shRNA sequences are listed in Supplementary Table 1. The next day, the medium was replaced with fresh complete growth medium, and cells were incubated overnight. The following day, virus-containing supernatants were harvested, centrifuged at 1000 r.p.m. for 5 minutes, and filtered through 0.45 μm porous membranes (STARLAB). Lentiviral transductions were performed in a 6-well plate format (3 × 10^5^ cells/well) using 10 µg/mL polybrene (Merck Millipore). Stably transduced cells were flow-sorted.

### Luciferase reporter assays

A375 melanoma cells were transfected with a pEZX-PG04 plasmid expressing Gaussia luciferase (GLuc) under the influence of the *PARP14* promoter (GeneCopoeia). After 48 hours, 5000 cells were seeded in triplicate in 96-well plates and were treated with increasing concentrations of IFNγ for 24 hours. Subsequently, 100 µL of supernatant was collected for further analysis, and the plate was stained with crystal violet for normalisation. The luminescence assay was carried out using the Genecopoeia Secrete Pair™ Dual Luminescence Assay kit according to the manufacturer’s instructions. Luminescence was measured on a Biotek Synergy™ H1 Hybrid Multi-Mode Reader with normalisation to the crystal violet absorbance values.

### Quantitative real-time PCR (RT-PCR)

Total RNA was extracted using QIAzol Lysis Reagent and isolated using RNAeasy mini kit (both from QIAGEN). cDNA synthesis was performed using the ProtoScript II First Strand cDNA Synthesis Kit (NEB). Expression levels of target genes were determined by RT-PCR using SensiMix SYBR No-Rox (Bioline) with the primers shown in Supplementary Table 1. Reactions were run on a Stratagene MX3000P real-time thermal cycler (Agilent Technologies). Relative expression levels were calculated using the 2^−ΔCt^ method after normalising to the expression levels of the housekeeping gene, *Gapdh*. Fold change levels were calculated using the 2^−ΔΔCt^ method after normalising to the untreated control.

### Western blot

Total proteins were extracted using SDS lysis buffer (4% SDS; 20% glycerol; 0.004% bromophenol blue; 0.125M Tris-Cl, pH 6.8; 10% 2-mercaptoethanol) and sonication (50 kHz for 30 seconds; VibraCell X130PB, Sonics Materials) at 4°C and subsequently denatured at 95°C for 5 minutes. Proteins were separated on RunBlue 4-12% bis-tris polyacrylamide gels (Expedeon) and then transferred onto iBlot PVDF membranes (ThermoFisher) using the Wet/Tank Blotting Systems (Bio-Rad). Membranes were probed overnight at 4°C in blocking solution containing the primary antibody. Primary antibodies used in this work were PARP14 (C-1) (Santa Cruz Biotechnology, sc-377150), Phospho-Stat1 (pSTAT1; Tyr701; 58D6) (Cell Signalling Technology, 9167L), STAT1 (Cell Signalling Technology, 9172), MHC Class I H2 Kb (Abcam, ab93364), GAPDH (Proteintech, 60004-1-Ig), TAP1 (Cell Signalling Technology, 12341), TAP2 (Cell Signalling Technology, 12259; Santa Cruz Biotechnology, sc-515576), and PD-L1/B7-H1 (R&D Systems, AF1019-SP; Cell Signalling Technology, 13684. This was followed by incubation with the appropriate secondary antibody for 1.5 hours at room temperature. Signals were developed using the Clarity Max Western ECL blotting substrate (Bio-Rad) and visualised on a Gel Doc XR+ Gel Documentation System (Bio-Rad).

### RNA sequencing

Total RNA was isolated using the RNAeasy mini kit. RNA integrity was assessed on an Agilent 2200 TapeStation (Agilent Technologies). RNA samples (~1 μg) were submitted for RNA sequencing (100 nt paired-end reads, <30 million reads per sample) using an Illumina HiSeq4000. Three samples per condition were sequenced. The read quality was assessed using FastQC. Raw reads were trimmed using trimmomatic (version 0.36.6; sliding window trimming with 4 bases averaging and average quality minimum set to 20). Trimmed reads were aligned to the reference genomes hg38_analysisSet (human) or mm10 (mouse) using HISAT2 (version 2.1.0; default parameters). Aligned reads were counted against GENCODE release 25 (human) or GENCODE release M14 (mouse) using htseq-count (version 0.9.1). Differential gene expression analysis was performed using edgeR (version 3.24.1). Heatmap generation and clustering were performed using Multiple Experiment Viewer version 10.2.

### Gene set enrichment analysis (GSEA), pathway analysis, and gene ontology (GO) enrichment analysis

GSEA was carried out using Broad Institute GSEA 3.0 software^56^ and the Broad Institute Molecular Signatures Database (MSigDB) version 7.0 [http://software.broadinstitute.org/gsea/msigdb/index.jsp]. For the analysis, the “MaxMean” test statistic was used to test enrichment using a two-class comparison when comparing groups or quantitative analysis for continuous variables. Genes were ranked based on the signal-to-noise ratio. All *P*-values and false discovery rates (FDR) were based on 500-1,000 permutations. GSEA using BubbleGUM software was performed using “BubbleMap” settings^57^.

Pathway overrepresentation analysis was conducted using Ingenuity Pathway Analysis^58^ (QIAGEN) version 01-12 or REACTOME version 70^59^ [https://reactome.org]. Genes used for pathways were pre-filtered to remove lowly expressed genes. GO enrichment analysis was performed using GOrilla^60^ [http://cbl-gorilla.cs.technion.ac.il], using two unranked lists of genes (target list and corresponding background list) as running mode. Process, function, and component GO terms were analysed for enrichment. The significance threshold was set to *P* < 0.001. GO enrichment analysis data are shown as bar graphs of –log(*P*-value) or scatterplots created using REViGO^61^ [http://revigo.irb.hr].

### Publicly available transcriptomics data retrieval and analysis

We accessed one transcriptomic dataset from pre-treatment and on-treatment biopsies of melanoma patients undergoing ICBT^28^ [GSE91061; https://www.ncbi.nlm.nih.gov/geo/]. Fragments per kilobase of transcript per million mapped fragments (FPKM) and transcripts per million (TPM) data were obtained from the GEO database. TPM values were converted to log_2_(TPM+1).

### Tumour immune infiltrate analysis by flow cytometry

When tumours reached the required endpoint volume, mice were sacrificed by cervical dislocation, and tumours were dissected. Tumours were incubated for 45 minutes with 100 μg/mL of Liberase (Sigma-Aldrich) in serum-free media at 37°C and then pushed through a BD Falcon 100 µM nylon cell strainer using a syringe plunge. The cell suspension was centrifuged at 1300 r.p.m. and 4°C for 7 minutes, and cells were stained for 20 minutes protected from light with LIVE/DEAD™ Fixable Blue Dead Cell Stain Kit (ThermoFisher) diluted at 1:1000 in PBS. Subsequently, Fc receptors were blocked, and cells were stained with surface stain antibody mix for 45 minutes at 4°C in the dark. Cells were fixed and permeabilised using the Foxp3/Transcription Factor Staining Buffer Set (eBioscience) following the manufacturer’s instructions. Intracellular staining was performed for 45 minutes at 4°C in the dark, after which samples were analysed on a BD FACSymphony flow cytometer (BD Biosciences). For all antibodies, a non-stained cell sample and appropriate fluorescence minus one control were analysed as well. Data were analysed using FlowJo version 8.7.

### Antibodies and reagents for flow cytometry

The following antibodies or staining reagents were purchased from BioLegend: CD16/32 (clone 93), CD45 (clone 30-F11), CD3 (clone 145-2C11), CD4 (clone GK1.5), CD8α (clone 53-6.7), CD25 (clone PC61), CD62L (clone MEL-14), CD69 (clone H1.2F3), PD1 (clone 29F.1A12), CD44 (clone IM7), Granzyme B (clone QA16A02), LAG3 (clone C9B7W), TIM3 (clone RMT3-23), TCR γ/δ (clone GL3), NK1.1 (clone PK136), and KI67 (clone 16A8). FOXP3 (clone FJK-16s) and TCR beta (clone H57-597) were purchased from eBioscience.

### In silico immunophenotyping

We computationally investigated the relationship between PARP14 inhibition and tumour infiltration levels in mouse melanoma samples using our RNA sequencing data. Computational immunophenotyping was performed using xCell cell type enrichment analysis. The data were validated using xCell (n = 64)^26^. Statistical significance was determined by unpaired two-tailed Mann-Whitney test using GraphPad Prism (version 9.0).

## Statistical analysis

Statistical analysis was performed using GraphPad Prism (version 9.0) using different statistical tests as indicated in the figure legends. Effects with *P*-values < 0.05 were considered statistically significant.

## Data availability

All data generated and supporting the findings of this study are available within the paper. The RNA-seq data have been deposited in ArrayExpress with accession numbers E-MTAB-12194, E-MTAB-12195, and E-MTAB-12196. TCGA data used are publicly available at the Genomic Data Commons portal (https://portal.gdc.cancer.gov/). Source data are available for this paper. All other data supporting the findings of this study are available from the corresponding author upon reasonable request.

**Supplementary Table 1.**
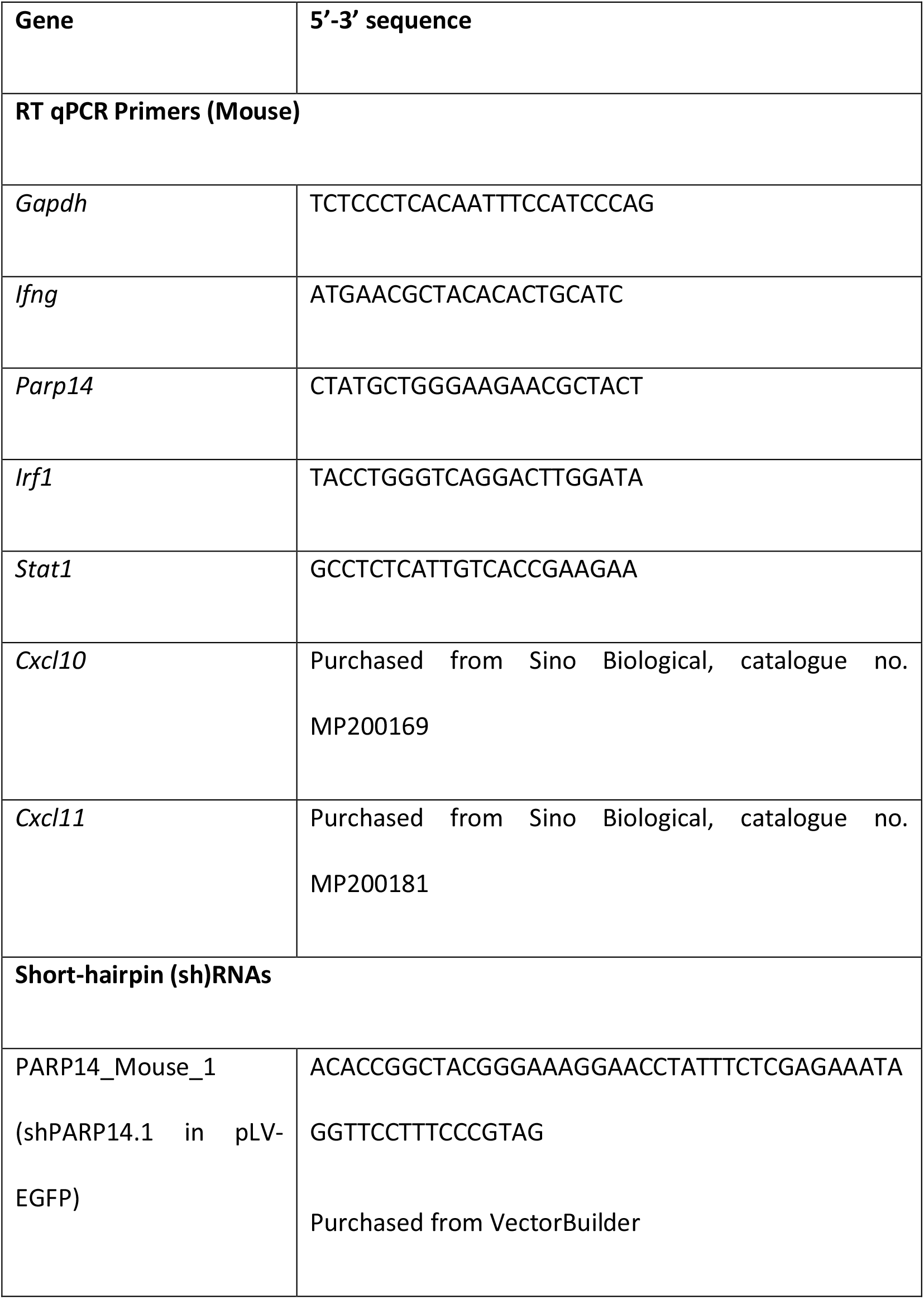
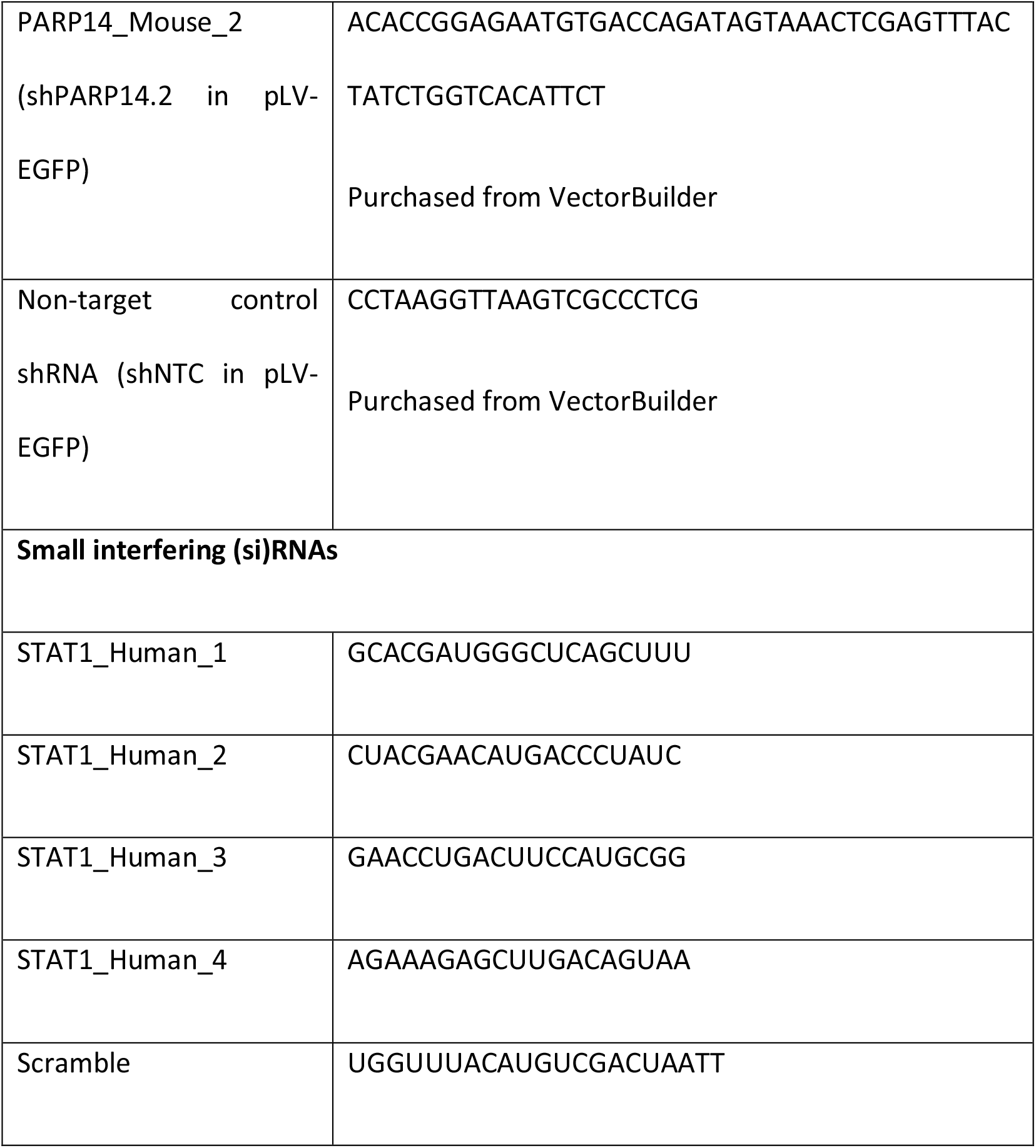
Oligonucleotides.

## Acknowledgements

This study was funded by a research sponsorship agreement between Ribon Therapeutics and The University of Manchester, grant 821034 from the Melanoma Research Alliance (MRA), grant 22-0091 from Worldwide Cancer Research (WCR), and awards from the Skin Cancer Research Fund (SCaRF) and Melanoma UK, all to A.H, and from the Christie Charity Fund to P.L. C.W.W. was funded by a scholarship from the Hong Kong Scholarship for Excellence Scheme (HKSES). C.E. was funded by a studentship from the Medical Research Council (MRC). We thank Chang Liu, Bin Gui, Dervla Isaac, and other employees of Ribon Therapeutics for helpful discussions and advice. We thank Dr Gareth Howell and The University of Manchester Flow Cytometry Core Facility for facilitating flow cytometry analysis, Dr Leo Zeef and the Genomics Technologies and Bioinformatics Core Facilities for assistance with RNA-seq, and members of the Biological Services Facility at The University of Manchester for help with animal work. We thank Michael Eisenstein for editing support. Some figures are created with BioRender.com.

## Author contributions

C.E., R.T.N, and A.H. conceived the study. C.W.W., C.E., K.N.S., R.L., K.S., M.L.F.C., E.U., H.M., B.A.T., K.K., N.R.P. and R.T.N. performed experiments. C.W.W., C.E., K.J.W., P.E.R., M.N., R.T.N., and A.H. contributed to experimental design and data analysis. A.H., D.J.W., M.P.S., K.J.W., P.E.R., R.T.N. and M.N provided supervision. C.W.W., C.E., K.N.S. and A.H. drafted the original version of this manuscript, with all authors reviewing subsequent drafts. P.L., M.N. and A.H. obtained funding for the study.

## Additional information

**Extended data** is available for this paper at…

**Supplementary information** The online version contains supplementary material available at…

**Correspondence and requests for materials** should be addressed to A.H.

