## Extended Data Figures for "PARP14 inhibition restores PD-1 immune checkpoint inhibitor response following IFNγ-driven adaptive resistance" (from another version of this preprint: version 1, 2022-11-18)

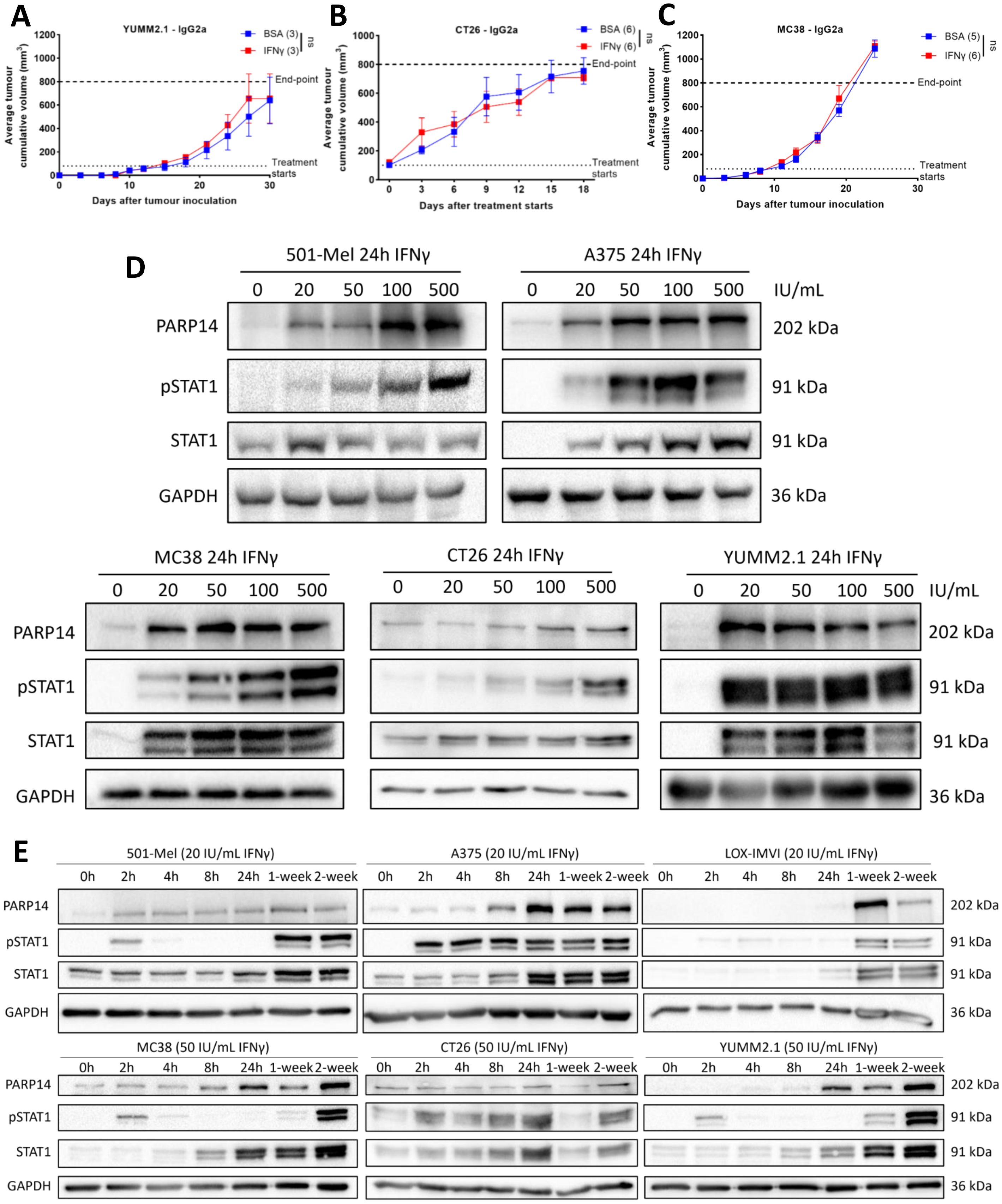

A
YUMM2.1 - IgG2a
BSA (3)
IFNγ (3)
ns
End-point
Treatment starts
Average tumour cumulative volume (mm3)
Days after tumour inoculation
B
CT26 - IgG2a
BSA (6)
IFNγ (6)
ns
End-point
Treatment starts
Days after treatment starts
C
MC38 - IgG2a
BSA (5)
IFNγ (6)
ns
End-point
Treatment starts
Days after tumour inoculation
D
501-Mel 24h IFNγ
A375 24h IFNγ
0 20 50 100 500 IU/mL
PARP14 202 kDa
pSTAT1 91 kDa
STAT1 91 kDa
GAPDH 36 kDa
MC38 24h IFNγ
CT26 24h IFNγ
YUMM2.1 24h IFNγ
E
501-Mel (20 IU/mL IFNγ)
A375 (20 IU/mL IFNγ)
LOX-IMVI (20 IU/mL IFNγ)
0h 2h 4h 8h 24h 1-week 2-week
MC38 (50 IU/mL IFNγ)
CT26 (50 IU/mL IFNγ)
YUMM2.1 (50 IU/mL IFNγ)

**Extended Data Fig. 1. Chronic IFNγ exposure does not impact growth rate in YUMM2.1/MC38 tumours; PARP14 and STAT1 activation are similarly upregulated in response to IFNγ.** (A-C) Data shown are from the experiment depicted in **Figure 1A**. Average tumour growth curve for (A) YUMM2.1, (B) CT26, and (C) MC38 cells after tumour inoculation or after treatment with the control IgG2a antibody. Statistical significance is determined at the last day of IgG2a treatment by Unpaired t test. NS, $P > 0.05$. (C) 501-Mel, A375, MC38, CT26, and YUMM.2.1 tumour cell lines treated for 24 hours with increasing concentrations of IFNγ (0, 20, 50, 100 and 500 IU/mL), with PARP14, pSTAT1 and STAT1 protein expression determined by western blot relative to a GAPDH loading control. (D) Treatment of human (501-Mel, A375, and LOX-IMVI) or mouse (MC38, CT26, and YUMM2.1) tumour cell lines with 20 IU/mL (human) or 50 IU/mL (mouse) IFNγ for up to 2 weeks. PARP14, pSTAT1 and STAT1 protein expression were determined via western blot relative to a GAPDH loading control.

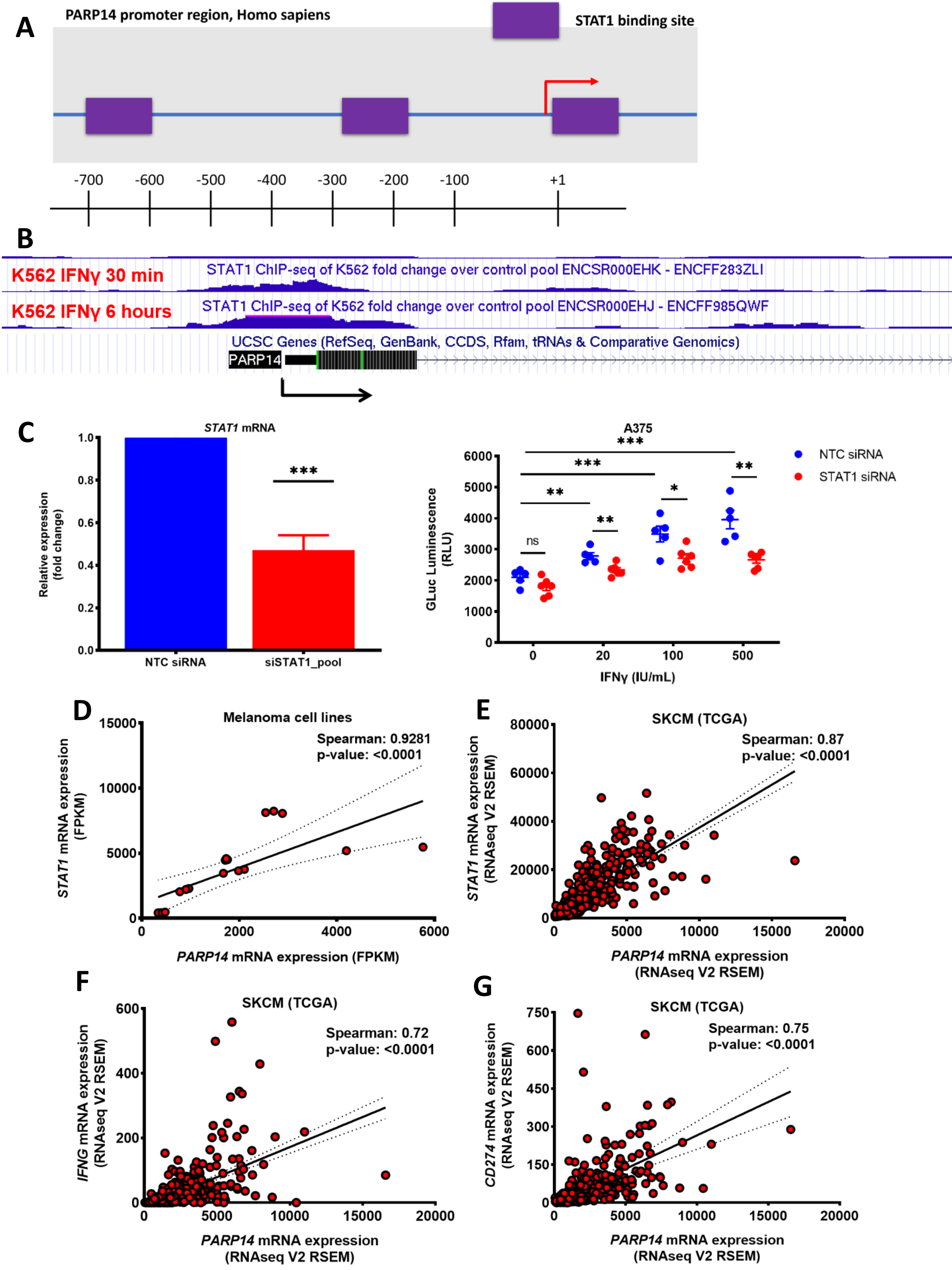
A
PARP14 promoter region, Homo sapiens
STAT1 binding site
-700 -600 -500 -400 -300 -200 -100 +1
B
K562 IFNγ 30 min
STAT1 ChIP-seq of K562 fold change over control pool ENCSR000EHK - ENCFF283ZLI
K562 IFNγ 6 hours
STAT1 ChIP-seq of K562 fold change over control pool ENCSR000EHJ - ENCFF985QWF
UCSC Genes (RefSeq, GenBank, CCDS, Rfam, tRNAs & Comparative Genomics)
PARP14
C
STAT1 mRNA
Relative expression (fold change)
NTC siRNA
siSTAT1_pool
***
A375
GLuc Luminescence (RLU)
IFNγ (IU/mL)
NTC siRNA
STAT1 siRNA
ns
D
Melanoma cell lines
Spearman: 0.9281
p-value: <0.0001
STAT1 mRNA expression (FPKM)
PARP14 mRNA expression (FPKM)
E
SKCM (TCGA)
Spearman: 0.87
p-value: <0.0001
STAT1 mRNA expression (RNAseq V2 RSEM)
PARP14 mRNA expression (RNAseq V2 RSEM)
F
SKCM (TCGA)
Spearman: 0.72
p-value: <0.0001
IFNG mRNA expression (RNAseq V2 RSEM)
PARP14 mRNA expression (RNAseq V2 RSEM)
G
SKCM (TCGA)
Spearman: 0.75
p-value: <0.0001
CD274 mRNA expression (RNAseq V2 RSEM)
PARP14 mRNA expression (RNAseq V2 RSEM)

**Extended Data Fig. 2. STAT1 is required for IFNγ-mediated PARP14 induction, and PARP14 and STAT1 expression strongly correlate in melanoma cell lines and patients.** (A) Schematic indicating positions of putative STAT1 binding sites (purple boxes) in the upstream PARP14 promoter region (~0.8 kb), as identified with MatInspector in silico analysis. The transcriptional start site is indicated by a red arrow. (B) ChIP-seq data analysis (data retrieved from ENCODE GSM935487 Snyder lab 2010) also verified the binding of STAT1 near the transcription start site of PARP14. (C) A375 cells were transfected with a GLuc reporter plasmid in which luciferase expression is controlled by the PARP14 promoter, and then treated with either a pool of siRNA targeting STAT1 or a NTC siRNA construct. Lefthand plot shows level of Stat1 expression upon treatment with each siRNA, righthand plot shows luciferase activity in response to 24-hour treatment with a range of IFNγ doses (0, 20, 100 and 500 IU/mL) in cells treated with either of the two siRNAs. Luminescence values are normalized to the cell confluency of each respective group, as determined by staining with crystal violet. Data are presented as mean ± S.E.M., with statistical significance determined by two-way ANOVA; NS $P > 0.05$, *$P < 0.05$, **$P < 0.01$, ***$P < 0.001$. (D–G) mRNA expression scatter plots showing co-expression of (D) *PARP14* with *STAT1* in melanoma cell lines or of *PARP14* with (E) *STAT1*, (F) *IFNG*, and (G) *CD274* in melanoma (SKCM, TCGA) patients. Spearman correlations and P-values are shown.

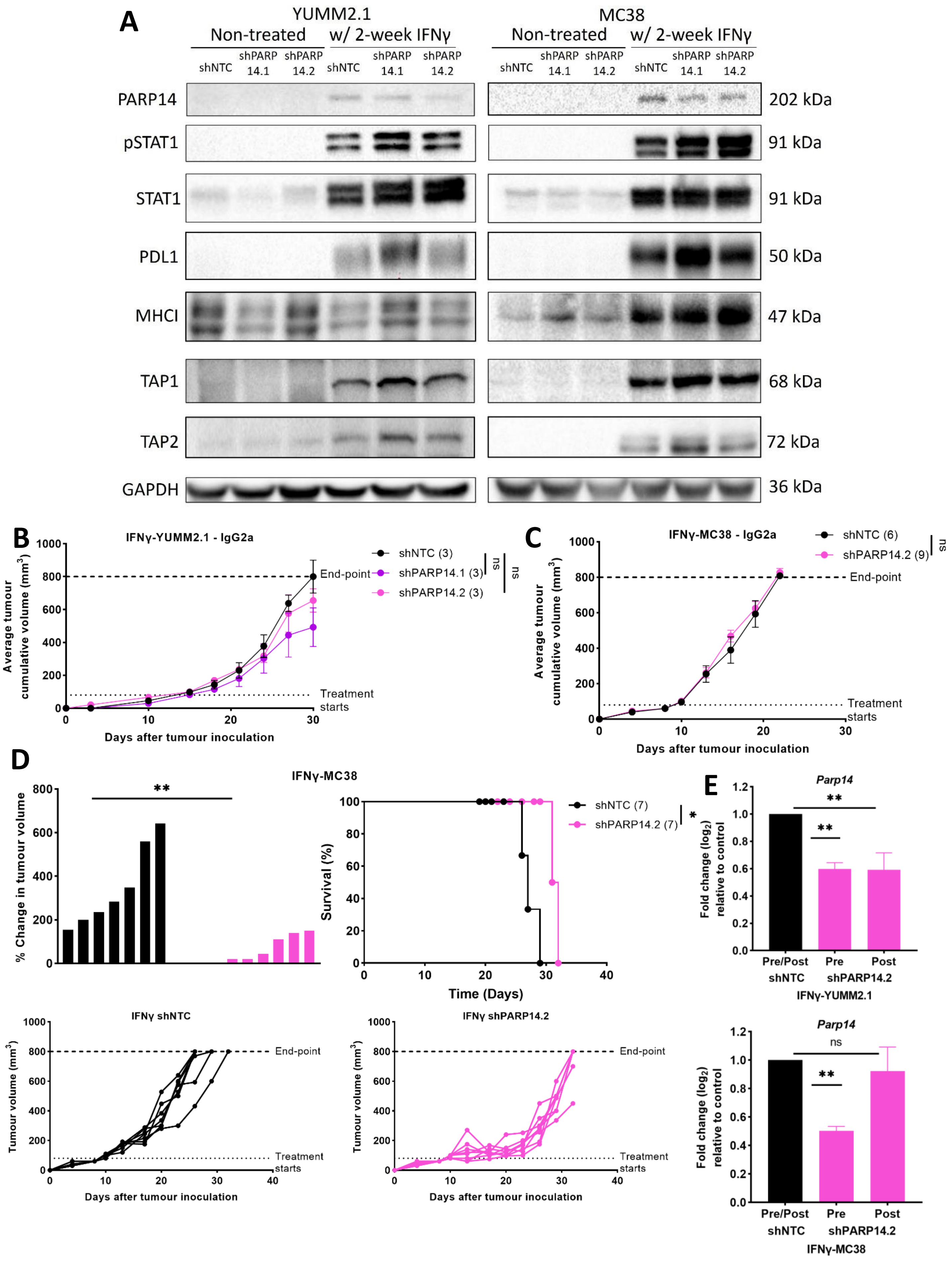

A
YUMM2.1
Non-treated
w/ 2-week IFNγ
MC38
shNTC
shPARP 14.1
shPARP 14.2
PARP14
pSTAT1
STAT1
PDL1
MHCI
TAP1
TAP2
GAPDH
202 kDa
91 kDa
91 kDa
50 kDa
47 kDa
68 kDa
72 kDa
36 kDa
B
IFNγ-YUMM2.1 - IgG2a
shNTC (3)
shPARP14.1 (3)
shPARP14.2 (3)
ns
ns
End-point
Treatment starts
Average tumour cumulative volume (mm$^3$)
Days after tumour inoculation
C
IFNγ-MC38 - IgG2a
shNTC (6)
shPARP14.2 (9)
ns
End-point
Treatment starts
Average tumour cumulative volume (mm$^3$)
Days after tumour inoculation
D
IFNγ-MC38
**
% Change in tumour volume
shNTC (7)
shPARP14.2 (7)
*
Survival (%)
Time (Days)
IFNγ shNTC
IFNγ shPARP14.2
Tumour volume (mm$^3$)
End-point
Treatment starts
Days after tumour inoculation
E
Parp14
**
**
ns
Fold change (log$_2$) relative to control
Pre/Post shNTC
Pre
Post
shPARP14.2
IFNγ-YUMM2.1
IFNγ-MC38

**Extended Data Fig 3. PARP14 ablation does not impact tumour growth rate but increases activation of STAT1 and expression of its target genes in chronic IFNγ-treated YUMM2.1/MC38.** (A) Western blot analysis of levels of PARP14, STAT1, and STAT1 targets in YUMM2.1 and MC38 cells with or without chronic IFNγ that were also treated with one of two PARP14 shRNAs or a negative control (shNTC). GAPDH is shown as a loading control. (B-C) Average cumulative tumour volume growth curves for (B) YUMM2.1 and (C) MC38 tumours, following the experimental protocol detailed in **Figure 2A**. Statistical significance determined at the last day of IgG2a treatment by Unpaired t test; NS $P > 0.05$. (D) Percentage of tumour volume change between the first dose of antibody treatment and administration of the final α-PD-1 dose (top left), corresponding Kalan-Meier plots (top right), and tumour growth rates (bottom) for the experimental protocol detailed in **Figure 2A**. Statistical significance determined by Unpaired t test or Log-rank (Mantel-Cox) test; *$P < 0.05$, **$P < 0.01$. (E) RT-qPCR analysis of *Parp14* mRNA expression in IFNγ-pre-treated and YUMM2.1 and MC38 tumours receiving shPARP14.2 or shNTC in pre- or post-tumour implantation. Data are presented as mean ± S.E.M. for six independent samples. Statistical significance determined by one-way ANOVA; NS $P > 0.05$, **$P < 0.01$.

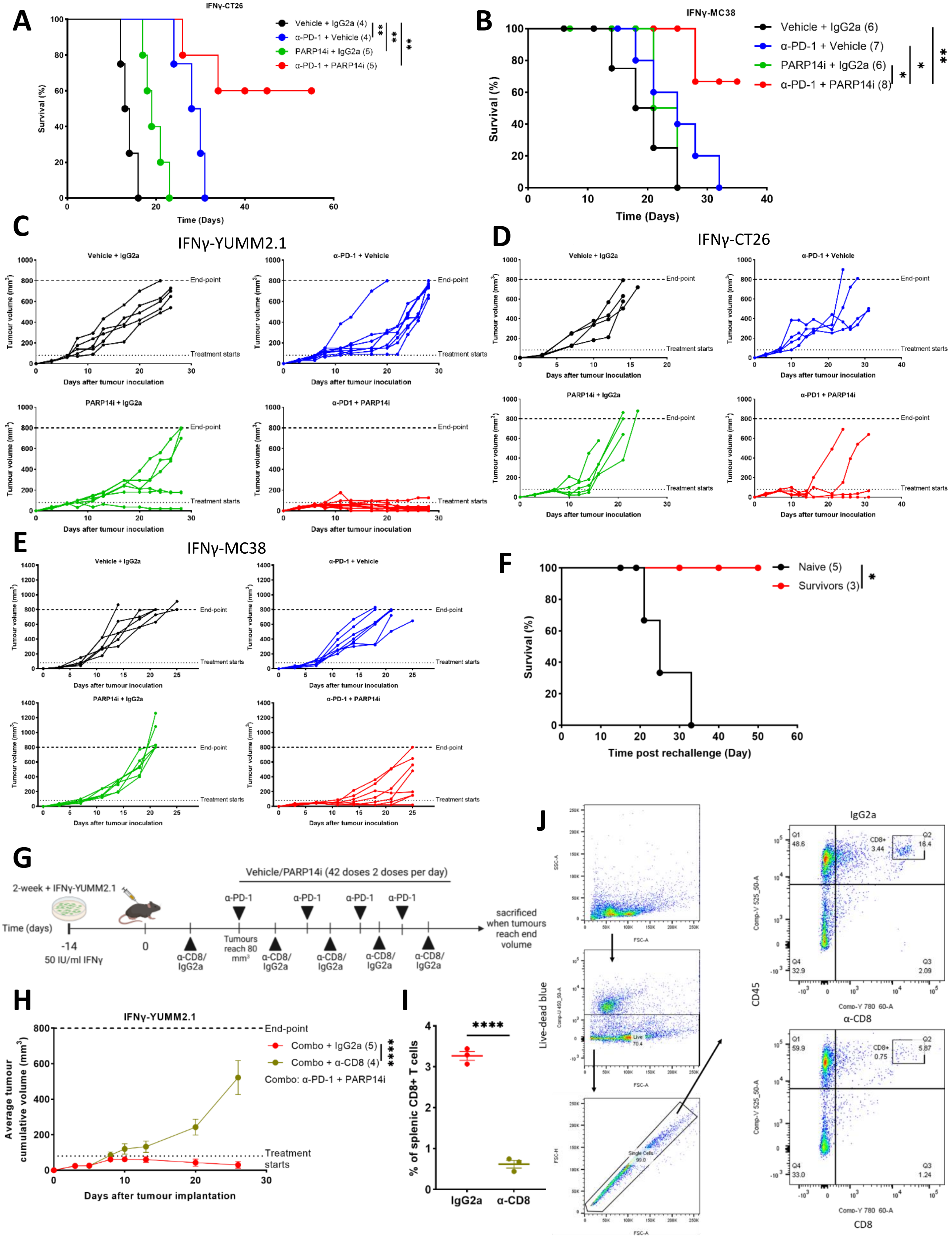
A
IFNγ-CT26
Vehicle + IgG2a (4)
α-PD-1 + Vehicle (4)
PARP14i + IgG2a (5)
α-PD-1 + PARP14i (5)
**
**
**
Survival (%)
Time (Days)
B
IFNγ-MC38
Vehicle + IgG2a (6)
α-PD-1 + Vehicle (7)
PARP14i + IgG2a (6)
α-PD-1 + PARP14i (8)
*
*
**
Survival (%)
Time (Days)
C
IFNγ-YUMM2.1
Vehicle + IgG2a
α-PD-1 + Vehicle
PARP14i + IgG2a
α-PD1 + PARP14i
End-point
Treatment starts
Tumour volume (mm³)
Days after tumour inoculation
D
IFNγ-CT26
Vehicle + IgG2a
α-PD-1 + Vehicle
PARP14i + IgG2a
α-PD1 + PARP14i
End-point
Treatment starts
Tumour volume (mm³)
Days after tumour inoculation
E
IFNγ-MC38
Vehicle + IgG2a
α-PD-1 + Vehicle
PARP14i + IgG2a
α-PD-1 + PARP14i
End-point
Treatment starts
Tumour volume (mm³)
Days after tumour inoculation
F
Naive (5)
Survivors (3)
*
Survival (%)
Time post rechallenge (Day)
G
Vehicle/PARP14i (42 doses 2 doses per day)
2-week + IFNγ-YUMM2.1
α-PD-1
Time (days)
-14
50 IU/ml IFNγ
0
α-CD8/ IgG2a
Tumours reach 80 mm³
sacrificed when tumours reach end volume
H
IFNγ-YUMM2.1
End-point
Combo + IgG2a (5)
Combo + α-CD8 (4)
****
Combo: α-PD-1 + PARP14i
Average tumour cumulative volume (mm³)
Treatment starts
Days after tumour implantation
I
****
% of splenic CD8+ T cells
IgG2a
α-CD8
J
SSC-A
FSC-A
Live-dead blue
Live
70.4
FSC-H
Single Cells
99.0
IgG2a
Q1
48.6
CD8+
3.44
Q2
16.4
Q4
32.9
Q3
2.09
Comp-V 525_50-A
Comp-Y 780 60-A
CD45
α-CD8
Q1
59.9
CD8+
0.75
Q2
5.87
Q4
33.0
Q3
1.24
CD8

**Extended Data Fig. 4. Pharmacological inhibition of PARP14 delays relapse of chronic IFNγ-treated CT26 and MC38 tumours when combined with α-PD1 therapy.** (A-B) Kalan-Meier plots of mice receiving IFNγ pre-treated CT26 (A) and MC38 (B) implants in the experiment shown in **Figure 2D**. Statistical significance determined by Log-rank (Mantel-Cox) test; *$P < 0.05$, **$P < 0.01$. (C-E) Growth rate for each tumour of IFNγ pre-treated YUMM2.1 (C), CT26 (D), MC38 (E) in the experiment shown in **Figure 2D**. (F) Survivors of α-PD-1 and PARP14i combinatorial therapy from the experimental protocol shown in **Figure 2D** were maintained for 60 days and then re-implanted with IFNγ pre-treated YUMM2.1 cells on the opposite flank to initial implant. Age-matched naïve mice act as a control group. The survival rate is shown for both groups. Statistical significance determined by Log-rank (Mantel-Cox) test; *$P < 0.05$. (G-J) Tumour-bearing mice received a total of four doses of α-PD-1 antibodies, 42 doses of PARP14i (twice daily for three weeks), and five doses of α-CD8 or IgG2a antibodies (a single dose every five days). (G) Experimental timeline. (H) Average cumulative tumour volume over the course of treatment. Statistical significance determined at day 24 of post tumour implantation by Unpaired t test; ****$P < 0.0001$. (I) Proportion of CD8+ cells among splenic T cells for animals receiving combination therapy with IgG2a (green) or α-CD8 (red). Statistical significance determined at day 24 of post tumour implantation by Unpaired t test; ****$P < 0.\ 0001$. (J) Flow cytometry gating strategy for assessing the efficiency of splenic CD8+ T cell depletion by α-CD8.

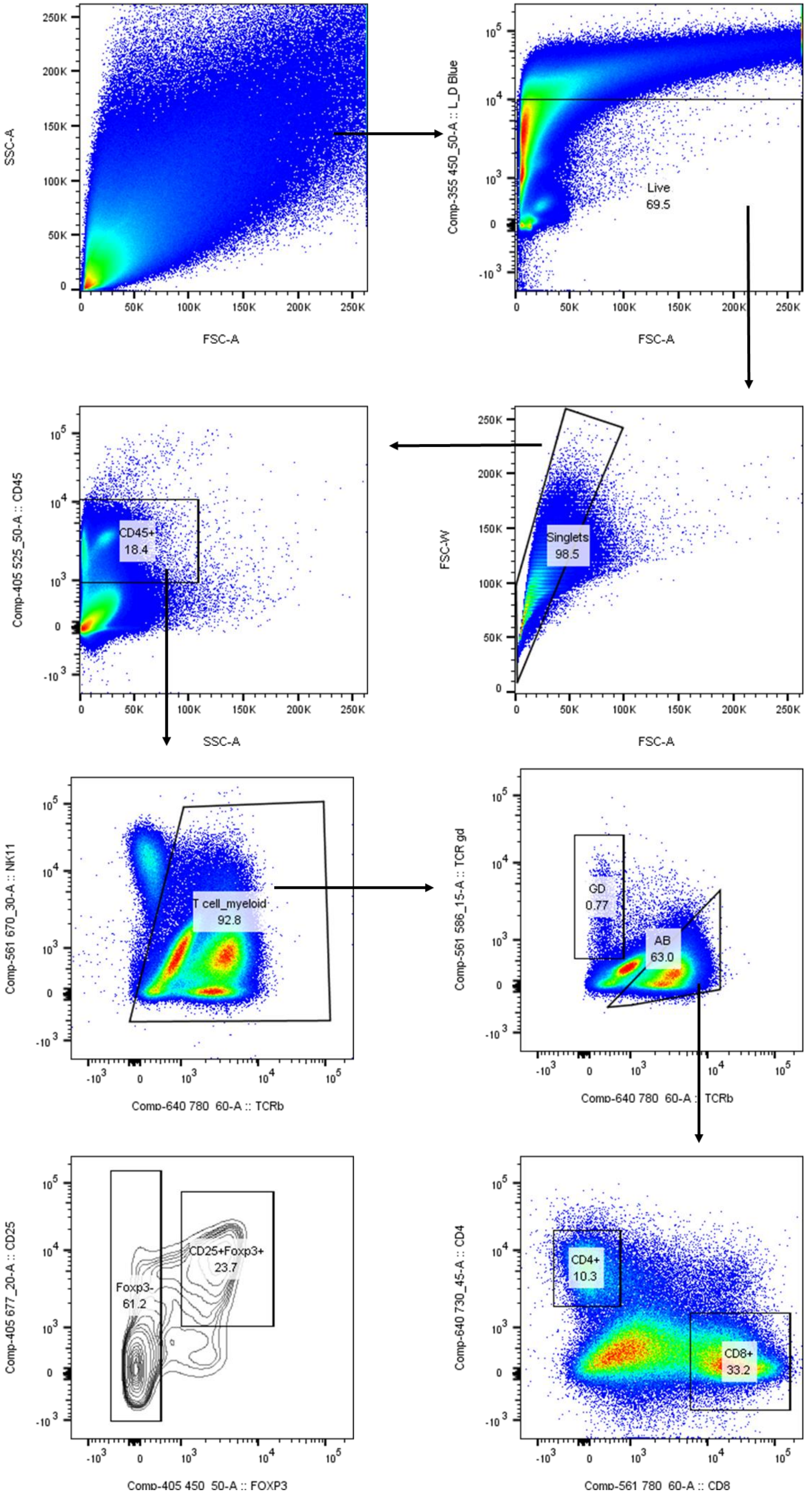

SSC-A
FSC-A
Comp-355 450_50-A :: L_D Blue
Live
69.5
FSC-W
Singlets
98.5
Comp-405 525_50-A :: CD45
CD45+
18.4
SSC-A
Comp-561 670_30-A :: NK11
T cell_myeloid
92.8
Comp-640 780_60-A :: TCRb
Comp-561 586_15-A :: TCR gd
GD
0.77
AB
63.0
Comp-640 780_60-A :: TCRb
Comp-405 677_20-A :: CD25
Foxp3-
61.2
CD25+Foxp3+
23.7
Comp-405 450_50-A :: FOXP3
Comp-640 730_45-A :: CD4
CD4+
10.3
CD8+
33.2
Comp-561 780_60-A :: CD8

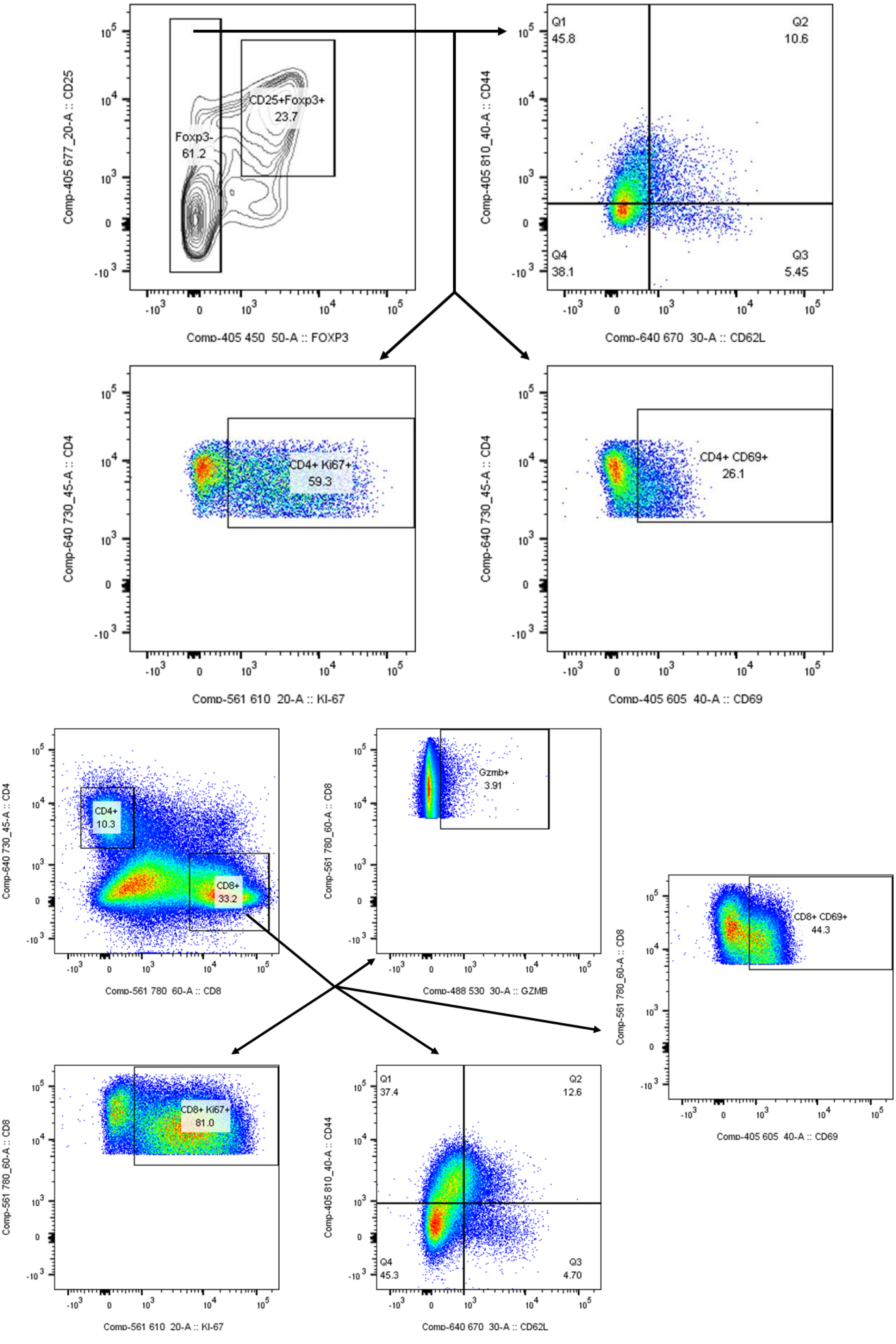

Comp-405 677_20-A :: CD25
Comp-405 450_50-A :: FOXP3
Foxp3-
61.2
CD25+Foxp3+
23.7
Comp-405 810_40-A :: CD44
Comp-640 670_30-A :: CD62L
Q1
45.8
Q2
10.6
Q4
38.1
Q3
5.45
Comp-640 730_45-A :: CD4
Comp-561 610_20-A :: KI-67
CD4+ Ki67+
59.3
Comp-405 605_40-A :: CD69
CD4+ CD69+
26.1
Comp-561 780_60-A :: CD8
CD4+
10.3
CD8+
33.2
Comp-488 530_30-A :: GZMB
Gzmb+
3.91
CD8+ CD69+
44.3
CD8+ Ki67+
81.0
Q1
37.4
Q2
12.6
Q4
45.3
Q3
4.70

**Extended Data Fig. 5. Flow cytometry gating strategy to assess the influence of sustained IFNγ exposure and PARP14 genetic ablation on tumour immune infiltrates.** Discrimination of leukocytes by forward scatter vs side scatter. Single cells were then selected using forward scatter width (FSC-W) vs forward scatter area (FSC-A) pulse geometry gating to discriminate between clumps and single cells. Live/dead cell-separation was achieved using LIVE/DEAD Fixable Blue Dead. Immune cells were then gated on CD45+ labeling. These can then be gated using the markers NK1.1, TCRβ, TCRγδ, CD4, CD8, and FOXP3 to quantify NK-T cells, NK cells, γδ T cells, TCRαβ$^{+}$ T cells, CD4$^{+}$ helper T cells, Tregs and CD8$^{+}$ T cells. To quantify activation status of these immune cell populations, we evaluated surface expression of CD69, CD44, and CD62L and intracellular expression of Ki-67 and GZMB.

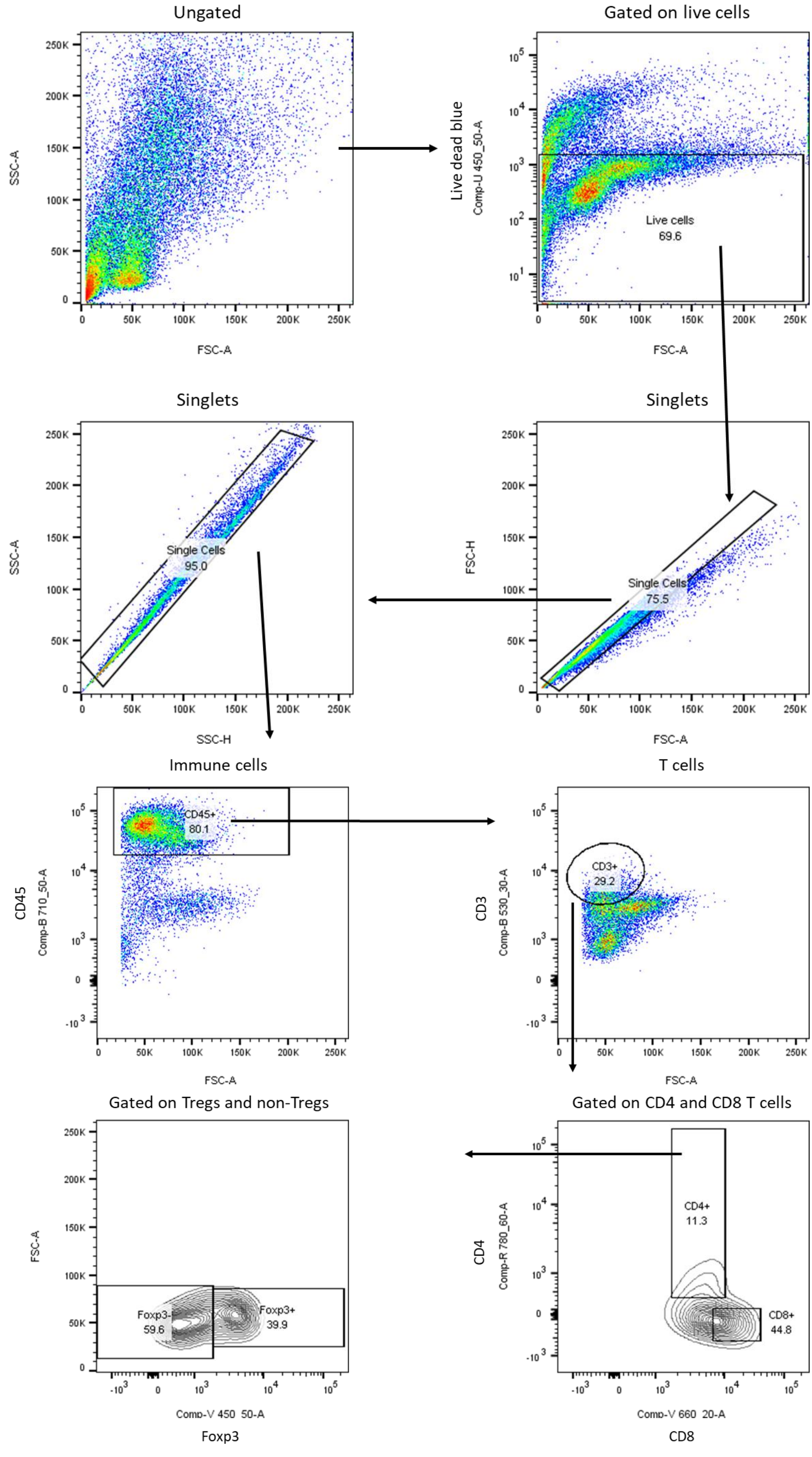
Ungated
Gated on live cells
SSC-A
FSC-A
Live dead blue
Comp-U 450_50-A
Live cells
69.6
Singlets
Singlets
SSC-A
SSC-H
FSC-H
FSC-A
Single Cells
95.0
Single Cells
75.5
Immune cells
T cells
CD45
Comp-B 710_50-A
CD45+
80.1
CD3
Comp-B 530_30-A
CD3+
29.2
Gated on Tregs and non-Tregs
Gated on CD4 and CD8 T cells
FSC-A
Foxp3-
59.6
Foxp3+
39.9
Comp-V 450_50-A
Foxp3
CD4
Comp-R 780_60-A
CD4+
11.3
CD8+
44.8
Comp-V 660_20-A
CD8

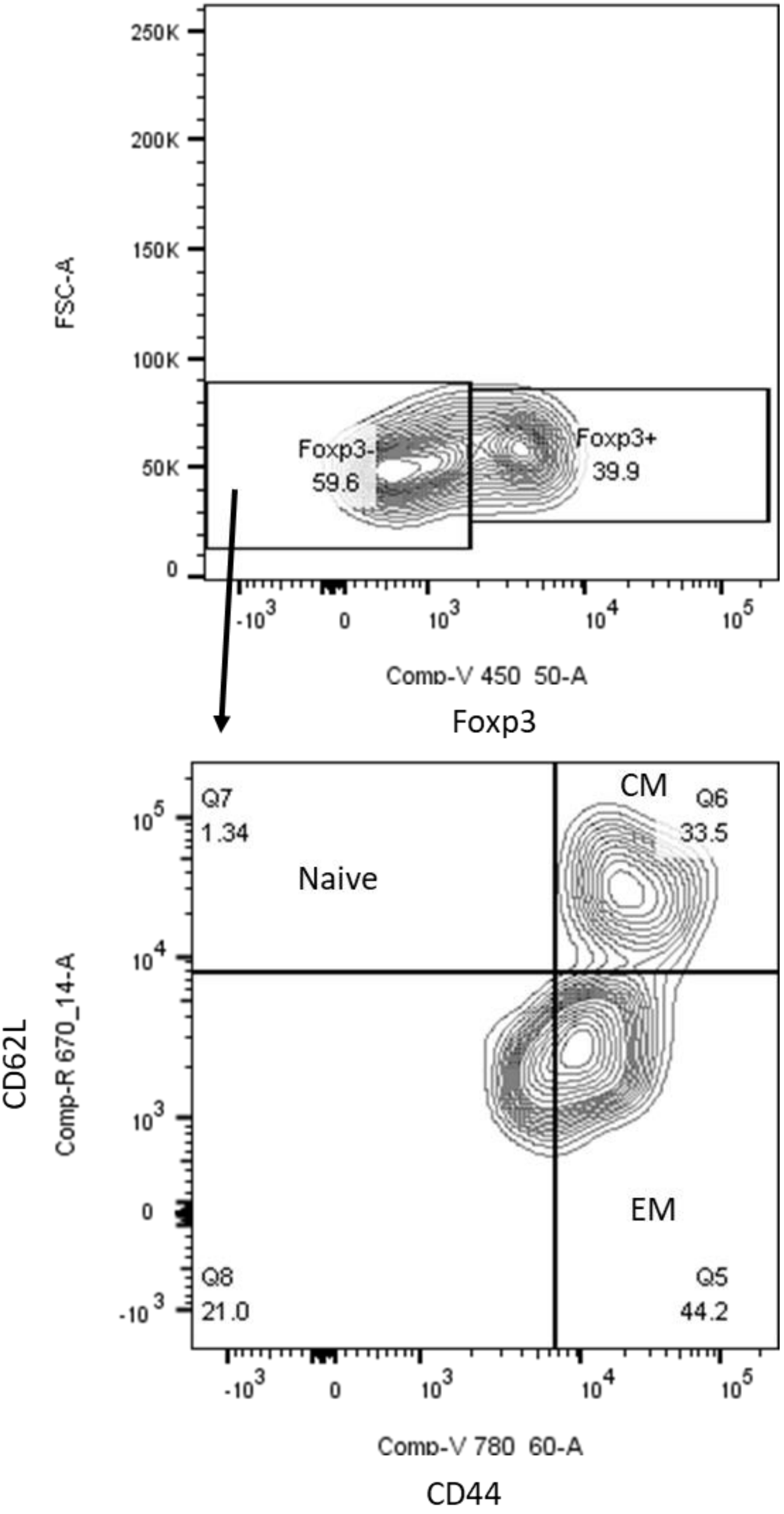

250K
200K
150K
100K
50K
0
FSC-A
Foxp3-
59.6
Foxp3+
39.9
-10^3
0
10^3
10^4
10^5
Comp-V 450_50-A
Foxp3
Q7
1.34
Naive
CM
Q6
33.5
10^5
10^4
10^3
0
-10^3
CD62L
Comp-R 670_14-A
EM
Q8
21.0
Q5
44.2
Comp-V 780_60-A
CD44

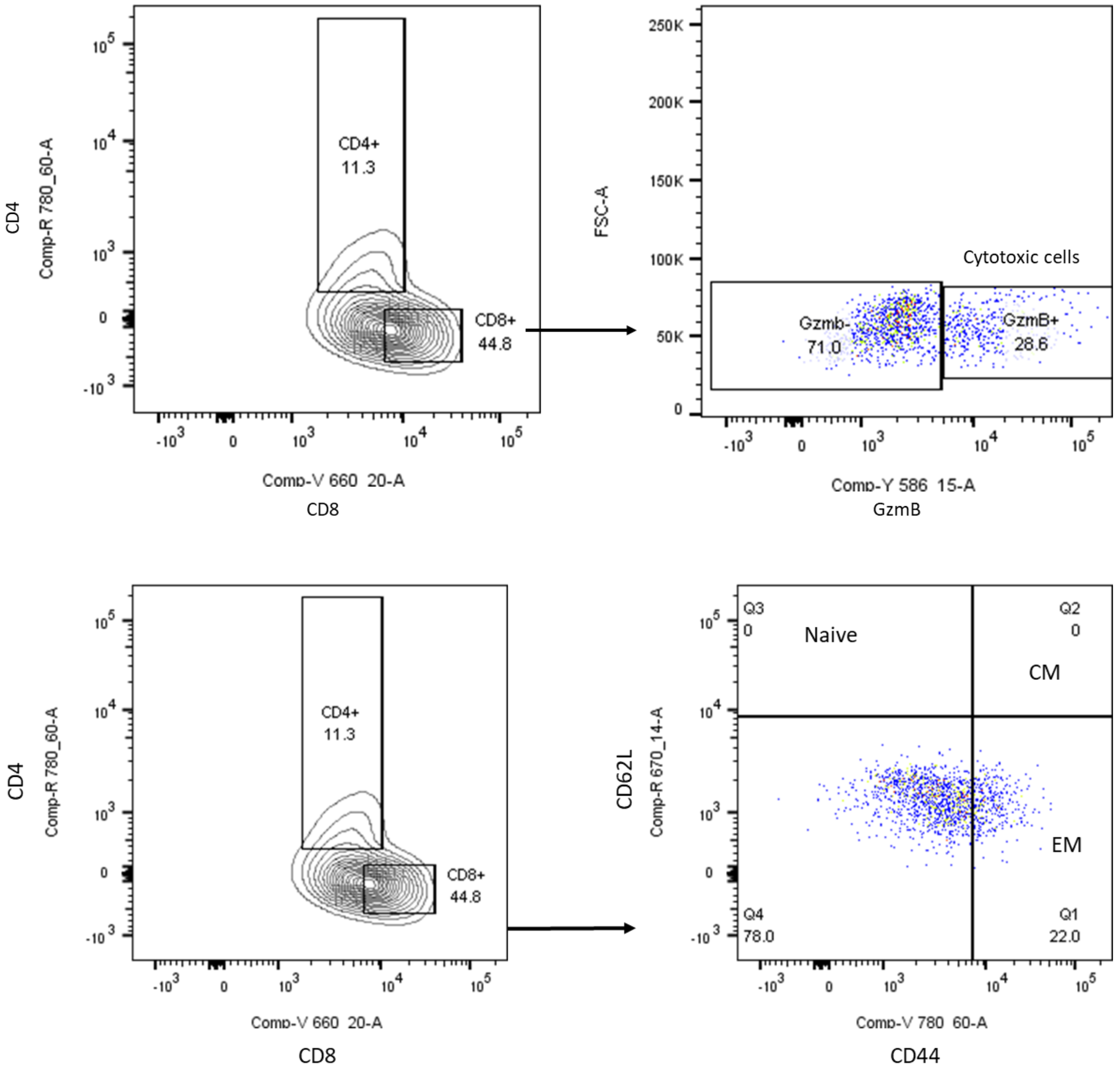
CD4
Comp-R 780_60-A
CD4+
11.3
CD8+
44.8
Comp-V 660_20-A
CD8
FSC-A
Cytotoxic cells
Gzmb-
71.0
GzmB+
28.6
Comp-Y 586_15-A
GzmB
CD4
Comp-R 780_60-A
CD4+
11.3
CD8+
44.8
Comp-V 660_20-A
CD8
Q3
0
Naive
Q2
0
CM
CD62L
Comp-R 670_14-A
EM
Q4
78.0
Q1
22.0
Comp-V 780_60-A
CD44

**Extended Data Fig. 6. Flow cytometry gating strategy for assessing the influence of PARP14 pharmacological inhibition on activation status of tumour-infiltrating immune cells.** Dead cells and debris were excluded, followed by single-cell gating to exclude duplets and cell clusters from further analysis. Immune cells gated by $CD45^{+}$ were then gated to quantify $CD4^{+}$ helper T cells, $T_{reg}$ cells and $CD8^{+}$ T cells, which we then analysed for surface expression and intracellular expression of CD69, CD44, CD62L, PD-1, LAG-3, TIM-3, GZMB.

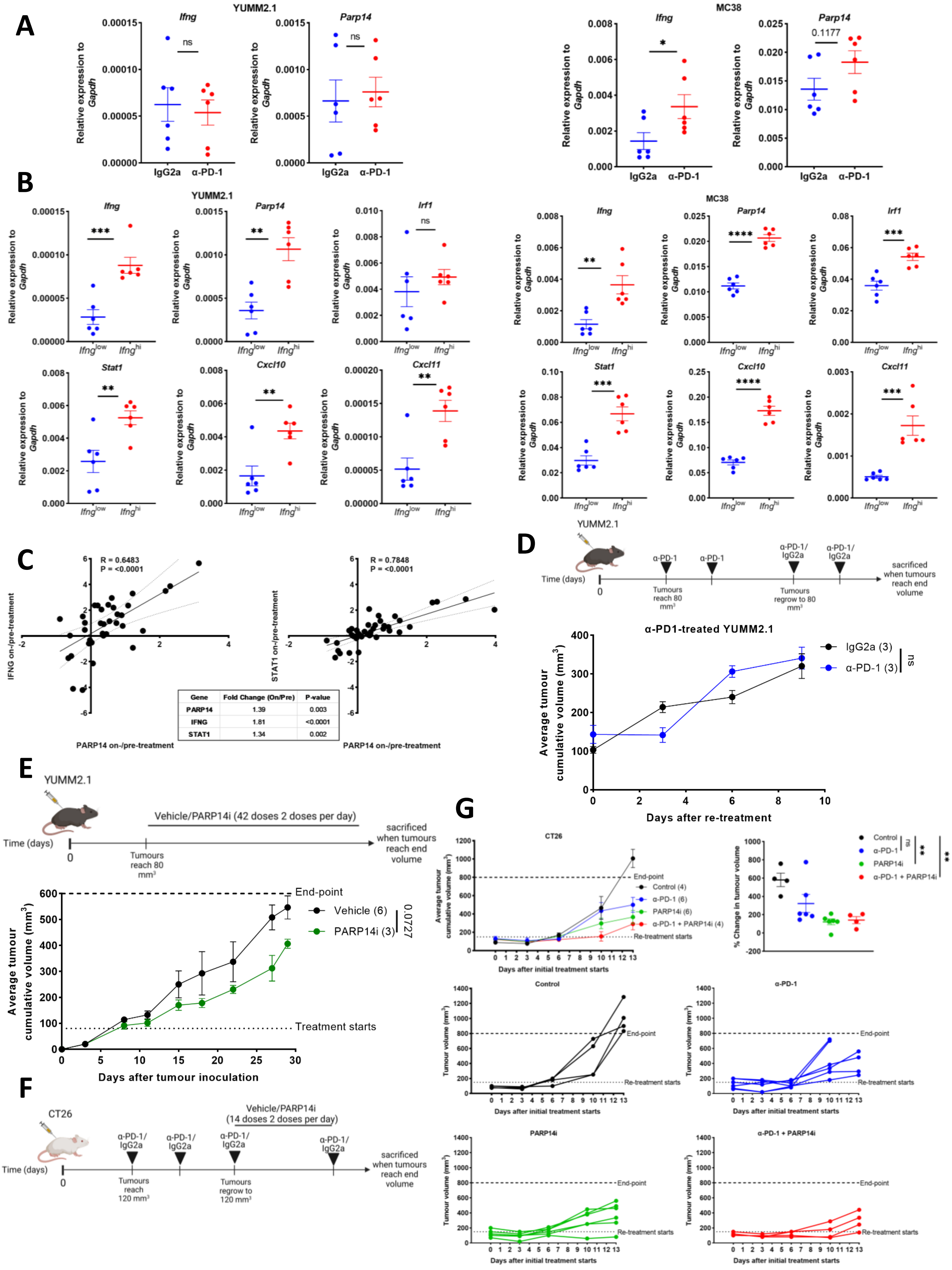
A
YUMM2.1
Ifng
Parp14
MC38
Relative expression to Gapdh
IgG2a
α-PD-1
ns
*
0.1177
B
YUMM2.1
MC38
Ifng
Parp14
Irf1
Stat1
Cxcl10
Cxcl11
Ifng low
Ifng hi
C
R = 0.6483
P = <0.0001
R = 0.7848
P = <0.0001
IFNG on-/pre-treatment
STAT1 on-/pre-treatment
PARP14 on-/pre-treatment
Gene | Fold Change (On/Pre) | P-value
PARP14 | 1.39 | 0.003
IFNG | 1.81 | <0.0001
STAT1 | 1.34 | 0.002
D
YUMM2.1
Time (days)
α-PD-1
α-PD-1/ IgG2a
Tumours reach 80 mm³
Tumours regrow to 80 mm³
sacrificed when tumours reach end volume
α-PD1-treated YUMM2.1
IgG2a (3)
α-PD-1 (3)
ns
Average tumour cumulative volume (mm³)
Days after re-treatment
E
YUMM2.1
Vehicle/PARP14i (42 doses 2 doses per day)
Tumours reach 80 mm³
End-point
Vehicle (6)
PARP14i (3)
0.0727
Treatment starts
Days after tumour inoculation
F
CT26
Vehicle/PARP14i (14 doses 2 doses per day)
Tumours reach 120 mm³
Tumours regrow to 120 mm³
G
CT26
Control (4)
α-PD-1 (6)
PARP14i (6)
α-PD-1 + PARP14i (4)
Re-treatment starts
Days after initial treatment starts
% Change in tumour volume
Control
α-PD-1
PARP14i
α-PD-1 + PARP14i
Tumour volume (mm³)

**Extended Data Fig. 7. PARP14 levels are augmented in tumours spontaneously relapsing after α-PD-1 treatment wherein it mediates resistance.** 8–12-week-old wild-type C57BL/6 mice were subcutaneously implanted with YUMM2.1/MC38 cells. Treatment with α-PD-1 or IgG2a antibodies (two doses, three days apart) was initiated once tumour volume reached ~100–150 $mm^3$. Within 48-hour of the last dose being administered, tumours were dissected, and the RNA was extracted and subjected to RT qPCR analysis. (A) *Ifng* and *Parp14* mRNA expression in mice receiving α-PD-1 or IgG2a. (B) YUMM2.1/MC38 αPD-1/IgG2a-treated tumours from the experiment described in panel A were reclassified as Ifnghigh and Ifnglow based on their expression levels. Expression levels for various STAT1 target genes (*Stat1*, *Irf1*, *Parp14*, *Cxcl10*, and *Cxcl11*) in these two cohorts are shown relative to the housekeeping gene *Gapdh*. Data are presented as mean ± S.E.M. for six independent samples. For A and B, significance was determined by unpaired t-test; NS $P > 0.05$, *$P < 0.05$, **$P < 0.01$, ***$P < 0.001$, ****$P < 0.0001$. (C) Correlation of fragments per kilobase of exon per million mapped fragments (FPKM) of *PARP14* versus *IFNG* and *STAT1* mRNA in matching melanoma patient biopsies on-treatment versus pre-treatment with anti-PD1 (n = 49, data retrieved from Riaz et al., 2017 cohort[27]). The mean is shown (black solid line) with a 99% confidence interval (black dashed line). Spearman correlations and associated P-values are shown. (D) 8–12-week-old wild-type C57BL/6 mice were subcutaneously implanted with YUMM2.1 cells. Treatment with α-PD-1 antibodies (two doses, three days apart) was initiated once tumour volume reached ~80 $mm^3$. Once the tumours had regrown to roughly the size at which α-PD-1 treatment was initially commenced, they were treated with two doses of IgG2a or α-PD-1 (three days apart). The average cumulative tumour volume growth curve is shown for each treatment condition. (E) YUMM2.1 cells were subcutaneously implanted into 8–12 weeks-old wild-type C57BL/6 female mice. Treatment of tumour-bearing mice was initiated once tumour volume reached ~80 $mm^3$. The cumulative tumour volume growth curve is shown for each treatment condition. Significance was determined on day 9 of re-treatment (D) and day 29 of post-tumour implantation (E) by unpaired t-test. NS $P > 0.05$. (F) 8–12-week-old wild-type BALB/C mice were subcutaneously implanted with CT26 cells. α-PD-1 antibody treatment (two doses, three days apart) was initiated once tumour volume reached ~120 $mm^3$. Tumours regrew and upon reaching 120 $mm^3$, the tumours were treated with two doses of α-PD-1 antibodies (1 dose every 3 days), two daily doses of PARP14i , vehicle or the combination therapy for a week. (G) Graphs show average tumour cumulative volume, percentage of tumour volume change between the start of α-PD-1 treatment and the end of re-treatment, and growth curves for each tumour. Percentage of tumour volume change statistical significance determined by one-way ANOVA; NS $P > 0.05$, **$P < 0.01$.
